# In depth analysis of kinase cross screening data to identify CAMKK2 inhibitory scaffolds

**DOI:** 10.1101/2020.01.08.883009

**Authors:** Sean N. O’Byrne, John W. Scott, Joseph R. Pilotte, André de S. Santiago, Christopher G. Langendorf, Jonathan S. Oakhill, Benjamin J. Eduful, Rafael M. Couñago, Carrow I. Wells, William J. Zuercher, Timothy M. Willson, David H. Drewry

## Abstract

The calcium/calmodulin-dependent protein kinase kinase 2 (CAMKK2) plays a central role in many cell signaling pathways. CAMKK2 activates CAMK1, CAMK4, AMPK, and AKT leading to numerous physiological responses. Deregulation of CAMKK2 is linked to several diseases, suggesting utility of CAMKK2 inhibitors for oncological, metabolic and inflammatory indications. In this work we review the role of CAMKK2 in biology and disease. Through analysis of literature and public databases we have identified starting points for CAMKK2 inhibitor medicinal chemistry campaigns. These starting points provide an opportunity for the development of selective CAMKK2 inhibitors and will lead to tools that delineate the roles of this kinase in disease biology.

## 1. Introduction

Calcium ions (Ca^2+^) are secondary messengers with important roles in cell signaling. Calmodulin (CaM) is a small protein that binds calcium ions *via* four EF-hand motifs.[1] Calcium bound calmodulin undergoes conformational changes, forming a complex with increased affinity for many CaM-binding proteins.[2-4] Experiments demonstrate that calmodulin binds to a broad range of enzymatic and non-enzymatic proteins.[5] A growing body of structural data suggests that although there are structural themes for CaM binding sites on proteins, there is substantial diversity in the binding sites and mechanisms by which the calcium-calmodulin complex influences protein structure and function.[6] Interestingly, a whole set of kinases has been given the classification of CAMK kinases.[7, 8] It turns out, however, that not all members of the CAMK group are regulated by interaction with CaM, and there are kinases outside the CAMK group with activity modulated by CaM.[9] Our group is interested in providing tool molecules that will allow the community to elucidate the many roles kinases play in disease biology. One particularly exciting CaM regulated kinase is CAMKK2, a serine/threonine kinase classified in the “other” subfamily of protein kinases.

CAMKK2 functions as a molecular hub to regulate critical cell functions. Once activated by binding to Ca^2+^/CaM, CAMKK2 phosphorylates numerous substrates that include the kinases CAMK1, CAMK4, and the alpha subunit of AMP-activated protein kinase (AMPK). Like CAMKK2, CAMK1 and CAMK4 are both activated by CaM binding, but for full activation they require phosphorylation by CAMKKs on their activation loops at Thr177 and Thr196 respectively.[10] CAMKK2 phosphorylates AMPK at α-Thr172, an event stimulated by binding of AMP or ADP to the AMPK regulatory γ subunit.[11] These three substrates influence cell functions such as cytoskeletal remodeling, cell cycle, motility, and inflammation (CAMK1); cell survival, gene expression, mRNA splicing, and immune response (CAMK4); and energy homeostasis, cytoskeleton remodeling, autophagy, and inflammation (AMPK) (**Figure 1**).[12, 13] The important oncology target AKT (also known as protein kinase B or PKB) has been established as an additional direct CAMKK2 substrate.[14-17]. AKT activation leads to phosphorylation of many substrates and a variety of well described cellular consequences, some of which are outlined in **Figure 1**.

**Figure 1:**
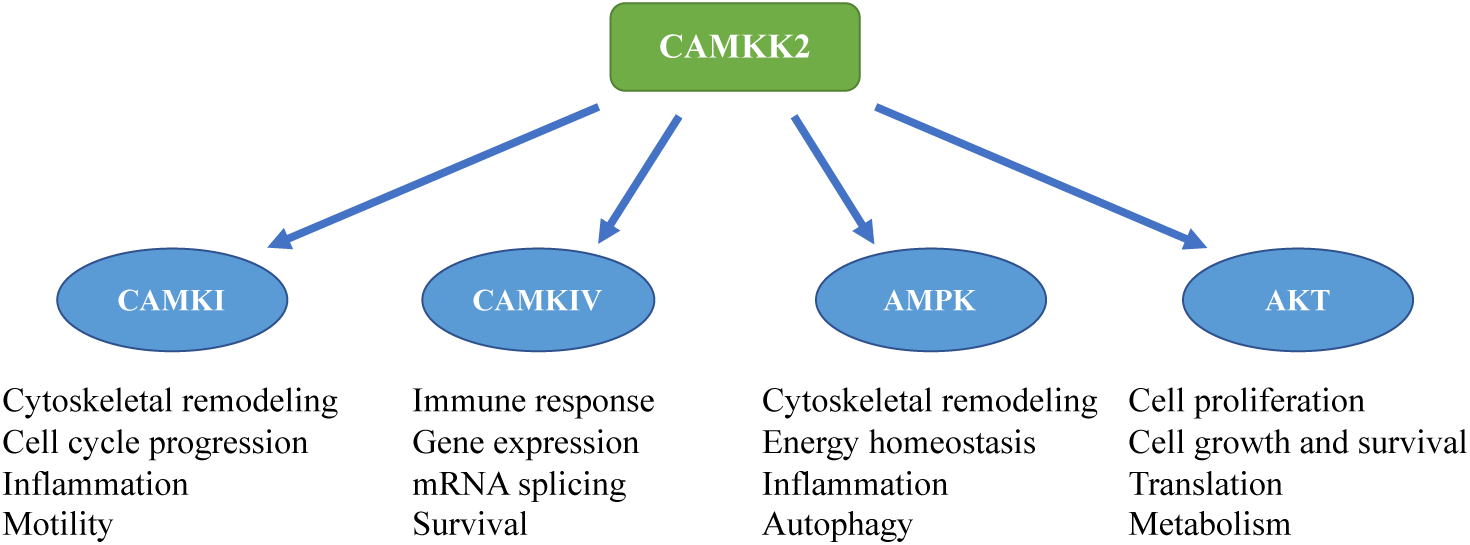
The CAMKK2 signaling pathway. Activation of CAMKK2 mediates downstream signaling of CAMK1, CAMK4, AMPK, and AKT.[18, 19]

This CAMKK2 signaling cascade regulates several distinct and important biological responses, including memory formation, cell proliferation, apoptosis and metabolic homeostasis.[20] CAMK1 is broadly expressed at low levels, and found at high levels in several areas of the brain.[21, 22] Evidence suggests that CAMK1 is involved in long term memory formation, regulation of cell growth, osteoclast differentiation and several other cellular functions.[23-29] CAMK4 is also mainly expressed in the brain, however it is also found in immune cells, the thymus, testes and ovaries.[30-33] CAMK4 is involved in cell proliferation, immunological and inflammatory responses, neurite outgrowth, memory formation and in the regulation of homeostatic plasticity.[34-39] AMPK is expressed ubiquitously and is involved in cell growth, metabolism and transcription, and plays a key role in the regulation of cellular and whole-body energy homeostasis.[40-42]. The outputs of AKT signaling have been well reviewed.[43-46]

Due to the importance of these signaling pathways, our group is interested in developing chemical probes for CAMKK2 to help elucidate the roles it plays in disease biology. Herein we provide a brief overview of CAMKK2 activation and structural biology, highlight literature that suggests the potential utility of CAMKK2 inhibitors, and outline chemotypes with CAMKK2 activity as potential starting points for drug discovery. Although CAMKK2 has not been the target of many medicinal chemistry campaigns, CAMKK2 inhibitory scaffolds can be found by careful analysis of broad kinome screening data in the literature. We highlight these CAMKK2 inhibitory scaffolds in order to provide useful information that will aid in the development of CAMKK2 selective chemical probes. These literature findings originate from a variety of different assay formats, we also acquired the compounds and assessed binding to CAMKK1 and CAMKK2 using differential scanning fluorimetry (DSF), and measured inhibition of CAMKK2 using a radiometric assay.

### 1.2 Structural features of CAMKK1 & CAMKK2

CAMKK1 and CAMKK2 are encoded by separate genes, both of which are subject to alternate splicing that generates multiple protein variants.[47, 48] CaMKK1 and CAMKK2 kinase domains have a high sequence homology and are structurally similar with an *N*-lobe consisting predominantly of *β*-sheets connected to an *α*-helix rich C-lobe *via* the ATP-binding hinge region. Like the other CaM kinases, CAMKK1 and 2 have a catalytic protein kinase domain adjacent to an autoinhibitory region that overlaps with the calmodulin-binding sequence.[49] Although CAMKK1 and -2 possess an autoinhibitory region, their biochemical properties are divergent. CAMKK1 explicitly requires calmodulin binding for effective activation, whereas CAMKK2 has substantial activity in the absence of the Ca^2+^-calmodulin (Ca^2+^-CaM) complex. The autonomous activity of CAMKK2 is regulated by several mechanisms including multi-site phosphorylation of a 23 amino acid region located N-terminal to the catalytic domain that we have termed the S3-node.[50] Cyclin-dependent kinase 5 (CDK5) phosphorylates S137 within the S3-node, which primes CAMKK2 for subsequent phosphorylation on S129 and S133 by glycogen synthase kinase 3β (GSK3β) and suppresses autonomous activity.[51-53] When bound to Ca^2+^-CaM, CAMKK2 undergoes autophosphorylation on T85, which generates autonomous activity by keeping CAMKK2 in the activated state after cessation of the Ca^2+^-stimulus.[54-56] Phosphorylation of CAMKK1 and 2 by cAMP-dependent protein kinase (PKA) leads to inhibition of Ca^2+^-CaM dependent activity, but not autonomous activity.[57]

There are crystal structures published for CAMKK1 complexed with hesperadin and GSK650394 (PBD: 6CCF & 6CD6).[58] CAMKK2 has 16 protein-inhibitor complexes published in the PDB.[59-61] The kinase domains of CAMKK1 and 2 are highly conserved and most inhibitors demonstrate activity towards both kinases. Despite this, there are some differences that could lead to preferential binding of inhibitors to either kinase as described by Santiago *et al.* in a recent publication.[58] They suggest two approaches that could be used to design specific inhibitors. First, there is a hydrophobic back-pocket near the kinase regulatory spine (R-spine) that consists of four residues which align to form a hydrophobic region when the kinase is in its active form. The R-spine contains residue Leu228 in CAMKK1 but is Met265 in CAMKK2. Type II inhibitors, which bind to the inactive DFG-out conformation of kinases can disrupt the R-spine and the Leu228-Met265 residue change could affect the flexibility and potential conformations of the R-spine. Hence inhibitors that disrupt the R-spine could utilize the differential movement to gain selectivity.[58]

The ATP-binding pockets of CAMKK1 and 2 have differences in residues that could also be exploited (**Figure 2**). Residues Leu233, Arg234 and Lys235 in CAMKK1 are replaced with Val270, Asn271 and Gln272 in CAMKK2. These changes cause the backbone to adopt a different conformation, primarily through the Leu233 in CAMKK1, which is larger than the equivalent Val270 in CAMKK2. Although challenging, the differences in residues/conformations in the hinge region could potentially be used to design selective inhibitors.[58]

**Figure 2:**
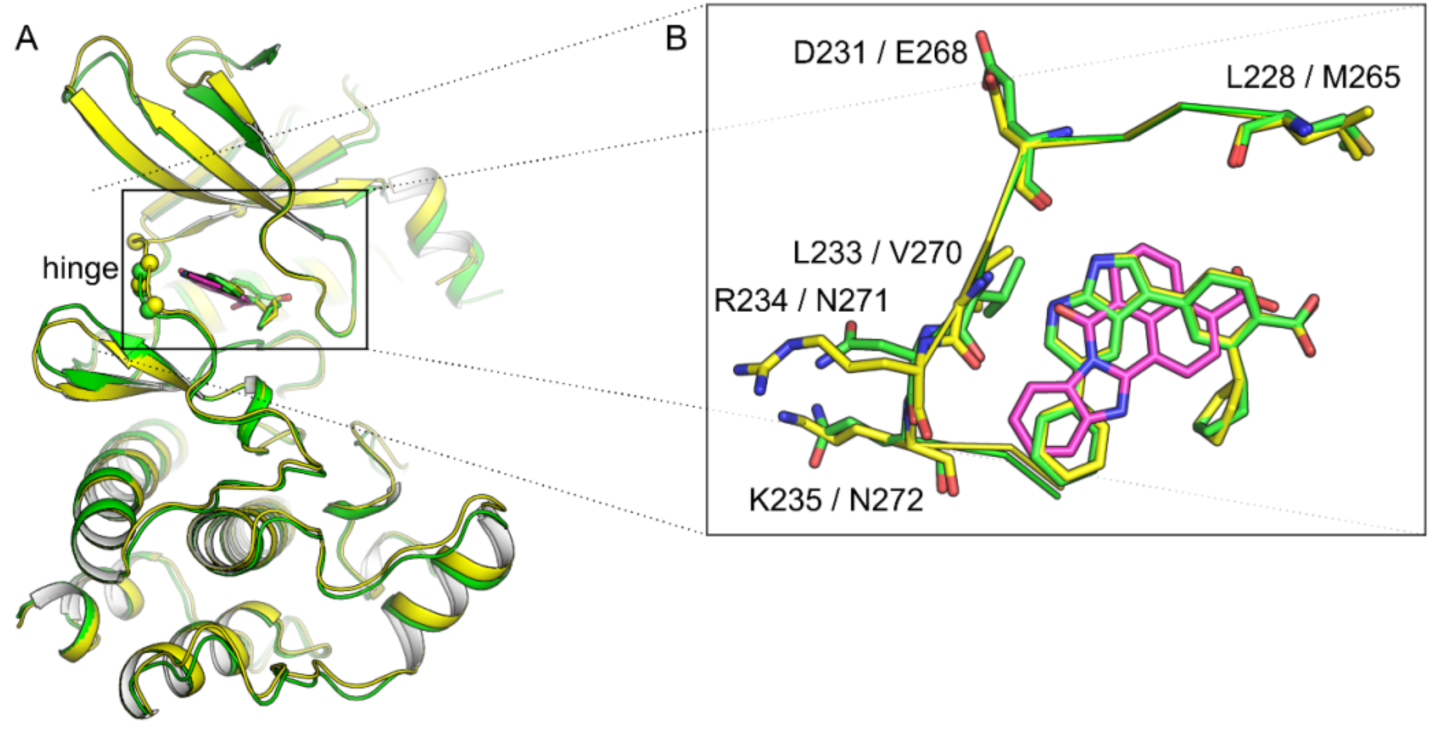
A) Superimposed structures of CAMKK1 (yellow, PDB 6CD6) and CAMKK2 (green, PDB 6BKU) bound to GSK650394. STO-609 (as bound to CAMKK2, PDB 2ZV2) is also shown in magenta. Spheres show positions where residues differ within the ATP-binding site of the two CAMKKs. B) Inset shows a top view of the ATP-binding sites of both CAMKKs.

### 1.3 CAMKK2 as a potential drug discovery target

In light of CAMKK2’s important role in several signaling pathways, it is no surprise aberrant activation can lead to improper biological functions. Inappropriate activation or overexpression of CAMKK2 has been implicated as a key driver in prostate, breast, ovarian, gastric and hepatic cancers.[62-70] Decreasing CAMKK2 activity using pharmacological inhibition or depletion of CAMKK2 using small interfering RNA (siRNA) reduces cell proliferation, migration and invasion in several cancer types.[68, 71-81]

The link between cancer and CAMKK2 is well documented in prostate cancer. In prostate cancer, the androgen receptor (AR)-signaling pathway is key in regulating disease progression, and is a major target for therapeutic intervention.[82, 83] Inhibition of the androgen receptor and reduction of circulating androgens have been the primary approaches used to treat the disease, but the therapies often succumb to drug resistance. CAMKK2 is upregulated in prostate cancer directly through the androgen receptor.[71, 78, 84] In cancer cell lines, CAMKK2 activates AMPK, altering cell metabolism and promoting cell proliferation.[78] There are seemingly conflicting reports on the role of CAMKK2 in LNCaP prostate cancer cell lines, indicating that further studies are required.[80, 84]. In spite of this, both downregulation with siRNA and pharmacological inhibition of CAMKK2 with the inhibitor STO-609 have been shown to reduce cell proliferation, migration, and invasion.[71, 78, 80, 84, 85]

Obesity, aberrant metabolism and type 2 diabetes are causative factors in non-alcoholic fatty liver disease (NAFLD), which is linked with rising incidence of hepatocellular carcinoma (HCC).[86, 87] CAMKK2 activation of hypothalamic AMPK stimulates ghrelin signaling, and promotes the desire to eat.[88] CAMKK2 inhibition has been shown to limit ghrelin-induced food intake in mice.[89] Furthermore, CAMKK2 null mice accumulate less body weight than wild-type mice when fed a high-fat diet, thereby offering protection against diet-induced obesity that promotes NAFLD.[90] In mouse models, treatment with STO-609 improved hepatic steatosis (fatty liver disease) compared to control groups.[91] CAMKK2 expression is significantly upregulated in HCC and correlates negatively with patient survival.[92] In several HCC cell lines, depletion of CAMKK2 using siRNA or inhibition of the enzyme with STO-609 decreased proliferation.[92] These studies suggest that CAMKK2 inhibition may be useful for the direct treatment of HCC, and also prevention of NAFLD, which is a risk factor for HCC.

In gastric cancer, studies using siRNA for knockdown and STO-609 for inhibition demonstrated decreased proliferation in various cell lines.[81, 93] Increased apoptosis was found in SNU-1 and N87 cancer cells, but not in normal gastric epithelial cells.[93] Increased expression of CAMKK2 in glioma correlates with negative outcome. Reducing CAMKK2 protein levels using siRNA leads to reduced proliferation, migration and invasion.[76] Similar results have been found in ovarian and breast cancer. Inhibition using STO-609 or siRNA leads to decreased proliferation and apoptosis in ovarian cancer cell lines, and to cell cycle arrest in the G1 phase in breast cancer cell lines.[68, 79] A recent publication demonstrated that CAMKK2 plays a key role in regulation of the immune-suppressive microenvironment in breast cancer.[94] CAMKK2 knockout mice had attenuated growth of grafted mammary tumors, which the authors attribute to a reduction in immunosuppressive activities. This phenotypic outcome was replicated using pharmacological inhibition with two chemically distinct CAMKK2 inhibitors.[95] A dominant negative CAMKK2 mutant showed reduced cell migration in DAOY medulloblastoma cell lines.[96] This study also investigated the effect of inhibition with STO-609 on migration, demonstrating reduced DAOY cell migration in a scratch assay.[96]

The potential therapeutic value of CAMKK2 inhibitors is not limited to the field of oncology. Due to the multitude of signaling pathways impacted by CAMKK2, and its role in whole-body energy homeostasis, this is of no surprise. Increasing evidence supports CAMKK2 inhibition as an avenue for the treatment of skeletal diseases. Osteoblasts (OBs) and osteoclasts (OCs) are key cells in bone tissue maintenance.[97] CAMKK2 is expressed in both OBs and OCs, and its inhibition increases osteoblast differentiation and bone growth, while suppressing osteoclast differentiation.[98] A subsequent study by the same authors provides evidence that CAMKK2 inhibition by STO-609 stimulated bone growth and reversed age-associated decline in bone strength, volume and several other parameters that are indicators of bone health.[99]

### 1.4 STO-609

The plethora of evidence discussed above indicates the promise of pharmacological intervention with CAMKK2 inhibitors. In the literature, pharmacological inhibition of CAMKK2 almost exclusively relies on the use of STO-609.[68, 71, 78, 93, 100-102] Although widely used, there is opportunity for improvement of this molecule. STO-609 is described as a selective inhibitor of the CAMKKs without significant activity on the other CaMKs. STO-609 is a potent inhibitor of CAMKK2 and has a five-fold lower K_i_ for CAMKK2 than CAMKK1.[103] A recent publication by York *et al*. characterized the metabolism, toxicity, pharmacokinetics, distribution and efficacy of STO-609.[104] Results in this paper demonstrate that STO-609 is well tolerated in mice without significant hepatic or renal toxicity. Exposure to human liver microsomes (HLM) and subsequent mass spectrometry analysis showed that STO-609 was metabolized to three mono-hydroxylated byproducts, principally by CYP1A2. In mice dosed intraperitoneally, the plasma half-life (t_1/2_) of STO-609 was between 8–12 hours. STO-609 was found in tissues that express CAMKK2, although low concentrations were found in skeletal muscle and brain.[103] The latter result is not unexpected as the physicochemical properties of STO-609 make it unlikely to cross the blood-brain barrier.

Although STO-609 has been widely used as a pharmacological inhibitor of CAMKK2 in the literature, it is not an ideal molecule, in part due to its poor aqueous solubility. STO-609 is a polycyclic aromatic compound containing five fused rings, and analysis of medicinal chemistry data sets suggests this is a liability.[105, 106] The extended planar ring system allows for significant π-π stacking, which leads to more stable crystal lattices, and contributes to poor solubility. Compounding this issue is the presence of the carboxylic acid moiety that also contributes to high crystal lattice energy *via* strong H-bond interactions. STO-609 is usually solubilized in DMSO or NaOH solution, which at higher doses can complicate use of the compound and lead to aberrant results. Indeed, a 10% DMSO solution is often used to kill cells *in vitro*, and as little as 0.25% DMSO can have inhibitory effects on some cell lines.[107] There are no published reports describing structure activity relationships for STO-609, or attempts to improve its properties. The available crystal structures of STO-609 bound to CAMKK2 could be used in attempts to improve STO-609 by adding solubilizing groups, increasing sp^3^ character, and decreasing the number of rings.

Another critical issue with STO-609 is its selectivity across the kinome. In the literature it is frequently described as being selective for CAMKK2. However careful examination revealed that STO-609 inhibits seven kinases with a similar potency to CaMKK1 (MNK1, CK2, AMPK, PIM2, PIM3, DYRK2, DYRK3).[108] Bain *et al* also found that at 1 μM, STO-609 was a more effective inhibitor of PIM3 than CAMKK2 and identified ERK8 as a collateral target. Further evidence to support these kinases as collateral targets of STO-609 is found in the MRC PPU (Medical Research Council Protein Phosphorylation Unit Dundee, UK) screening database where MNK1, PIM3, and ERK8 were inhibited by STO-609 at 1 μM.

Inhibitors having off-target activities can confound the mechanistic interpretation of phenotypic data, especially when the inhibitor in question is utilized at high concentrations in cellular or *in vivo* assays, which is frequently the case with STO-609. For example, the proto-oncogene PIM3 is a kinase that can prevent apoptosis and promote cellular survival, leading to tumorigenesis.[109] PIM3 has been implicated in hepatic, pancreatic, colon and gastric cancers.[110-113] The upregulation of CAMKK2 and PIM3 in HCC, and the fact that both are inhibited by STO-609, suggests that the results of some may not be solely due to CAMKK2 inhibition. It has also been shown that STO-609 binds and activates the aryl hydrocarbon receptor which could in some contexts be confounding due to induction of P450s or regulation of immune response.[114] The poor drug-like properties of STO-609 that require the use of high concentrations and non-ideal formulations coupled with its kinase off-target interactions underscore the need for a more selective inhibitor with better physiochemical properties that can be used to more accurately probe the functions of CAMKK2 in cells and *in vivo*.

## 2. Results

### 2.1 STO-609 – Selectivity

To further investigate the kinase selectivity profile of STO-609, it was screened across a panel of over 400 wild-type human kinases (KINOME*scan*®, DiscoverX). At 1 μM STO-609 was revealed to be a potent inhibitor of 13 kinases in addition to CAMKK2. The most potently inhibited kinases were CDKL2, GRK3, STK36, CSNK2A2, YSK4 and DAPK2. The kinome treespot and PoC values are shown in **Figures 3A** and **3B**. The full KINOME*scan* data is available in the SI (**Table S1**). These single concentration screening results require follow-up experiments in both dose-response and orthogonal assays to validate activity.

**Figure 3:**
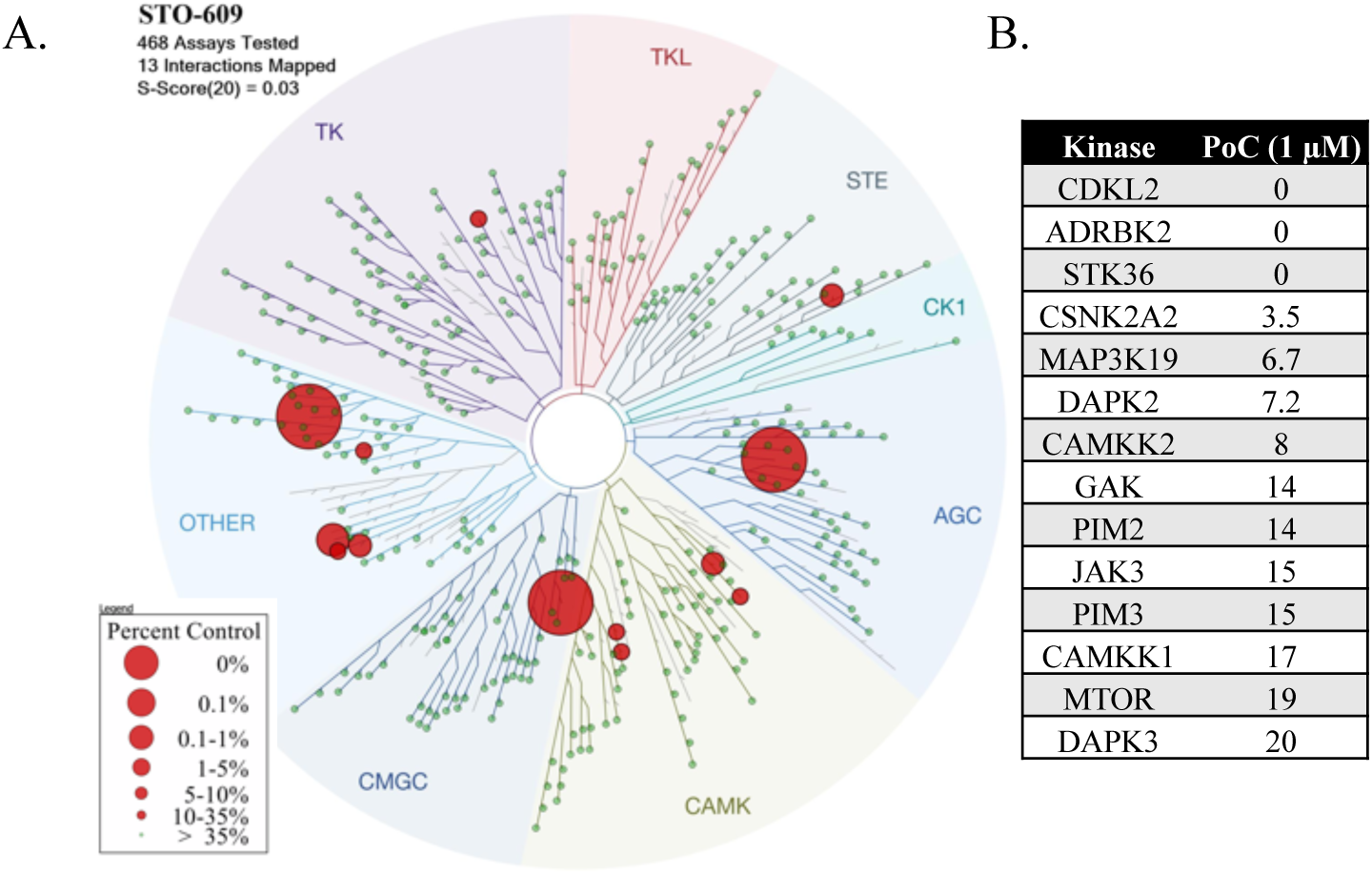
Selectivity and off-targets of STO-609 evaluated at 1 μM. **A**) Kinome treespot showing location of kinases that STO-609 binds to. **B**) List of kinases and their percent of control (PoC) remaining when treated with 1 μM of STO-609.

The KINOME*scan* results corroborate PIM2 and PIM3 as targets of STO-609. Bain *et al* report casein kinase 2 (CK2) as being potently inhibited with an IC_50_ of 190 nM.[115] In the KINOME*scan* data a truncated version of CK2, using only the catalytic subunit alpha2 (CSNK2A2), was also inhibited effectively following treatment with 1 μM of STO-609. CK2 is overexpressed in several cancers including breast, lung, prostate and kidney, and is associated with aggressive tumorigenesis.[116-120]

Three potential off targets identified in the KINOME*scan*, CDKL2, GRK3 and STK36, were not present in Bain *et al*’s or MRC’s profiling panels. CDKL2 and GRK3 have been implicated in breast cancer progression and their overexpression correlates with poor prognosis.[121, 122] STK36 is a node in the Hedgehog signaling pathway mediating GLI-dependent transcription.[123] These preliminary KINOME*scan* results require confirmation in orthogonal kinase inhibition assays.

The evidence of significant off-target kinase inhibition by STO-609 is compelling. Given that several of the kinases that are potently inhibited are also potential oncology targets, caution must be used when assigning experimental results exclusively to CAMKK2 inhibition. High quality probes are essential for elucidating the role of kinase signaling in healthy and diseased biological systems and it is possible that inaccurate conclusions may be drawn from use of non-selective probes.[124, 125] Ideally, concurrent testing of a chemically distinct CAMKK2 inhibitor should be used as an orthogonal probe to verify the mechanism of action.

### 2.2 Other CAMKK2 tool compounds

In the literature there are two recent publications that described potential CAMKK2 tool compounds. In the publication by Price *et al.* the inhibitors disclosed were based on 3,5-diaryl-7-azaindoles (**2**), 3,6-disubsituted-7-azaindoles (**3**), 2,4-diaryl-7-azaindoles (**4**) and 2-anilino-4-aryl-pyrimidine (**5**) cores.[89] In the publication by Asquith *et al*., 1,2,6-thiadiazin-4-ones (**6**) derivatives were shown to have only moderate CAMKK2 activity but a co-crystal structure was disclosed.[126] Structures of these compounds and STO-609 are depicted in **Figure 4**.

**Figure 4:**
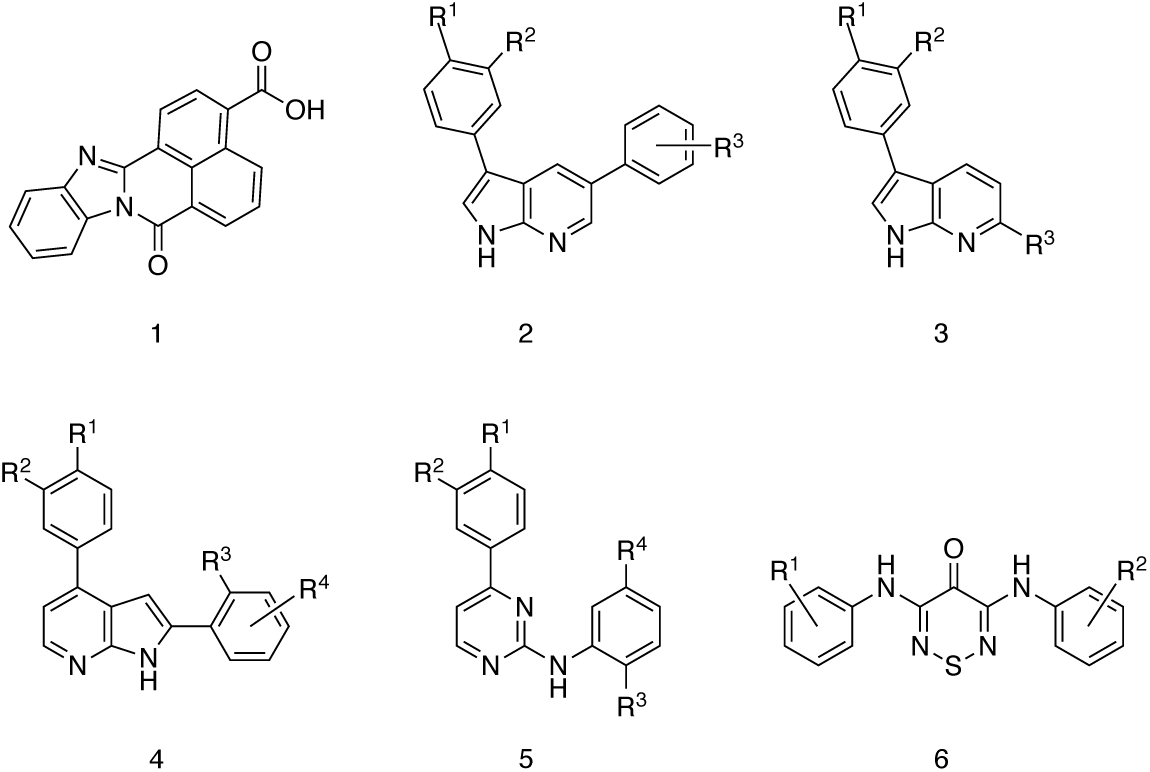
The most cited CAMKK2 inhibitor STO-609 (**1**). The scaffolds of CAMKK2 inhibitors (**2**-**6**) with structure activity studies described in the literature.[89, 126]

Price *et al* identified CAMKK2 inhibitors from a screen of 12,000 ATP-competitive kinase inhibitors and similar pharmacophores. The azaindole and anilino-pyrimidine cores are well documented as Type 1 kinase inhibitors that displace ATP from the hinge region of the enzymes.[127] Although significant effort was invested in each series, affording potent single digit nanomolar CAMKK2 inhibitors in biochemical assays, various factors such as poor cell permeability, low solubility, and poor *in vivo* absorption meant that most compounds were unsuitable for probing CAMKK2 activity *in vivo*. In addition, broad kinase screening data for the compounds was not available, so kinome-wide selectivity is unknown. The authors were successful in finding a central nervous system (CNS) penetrating compound. When orally dosed, rats treated with the compound showed a 40% reduction in ghrelin-induced food intake compared to a control group.[89]

The 1,2,6-thiadiazin-4-one derivatives identified by Asquith *et al.* represent an unusual class of hinge binders. The compounds were screened against a kinase panel representing the major branches of the kinome. CAMKK2 hits were identified using a thermal shift assay. Analogues were synthesized based on the best compound from the initial screen yielding inhibitors with low micromolar CAMKK2 inhibitory activity. The co-crystallization of CAMKK2 with one of the inhibitors showed that the thiadiazinone carbonyl and one of the anilino NH groups interacted with the hinge region.[126]

### 2.3 Literature survey of CAMKK2 inhibitor chemotypes

Given the modest selectivity of STO-609, and the lack of alternative high-quality tool compounds, a survey of the literature and several public databases was undertaken to identify alternative CAMKK2 inhibitor chemotypes. Candidate CAMKK2 inhibitors were acquired and their binding assessed in a DSF assay and CAMKK2 inhibition enzyme activity assay.

The databases mined in this study were the ‘Kinase Profiling Inhibitor Database’ provided by the University of Dundee, the ‘Kinase Inhibitor Resource’ provided by the Fox Chase Cancer Center and Reaction Biology Corporation, ‘KInhibition’ provided by Fred Hutchinson Cancer Research Center and the LINCS KINOME*scan* database provided by Harvard Medical School. Using a selection criterion of >25% inhibition at 1 µM, 25 compounds were identified. The inhibitors represent a diverse range of chemotypes including 2,4-dianilinopyrimidines, 2-anilino-4-arylpyrimidines, indolinones, pyrazolo-pyrimidines, fused pyrimidines, quinolines and *bis*-indoles. The set was supplemented with additional compounds that were described as CAMKK2 inhibitors in the literature, or where CAMKK2 had been noted as a collateral target during broad kinase screening activity. The set was further expanded to include commercially available analogues for some of the chemotypes, such as 2-anilinopyrimidines and oxindoles. A total of 52 compounds (**Table S2**) were evaluated against CAMKK1 and CAMKK2 in a DSF and enzyme inhibition assay.

### 2.4 Differential Scanning Fluorimetry & Enzymology

DSF is commonly employed to evaluate inhibitor binding and stabilization.[128, 129] The compounds ability to stabilize purified kinase domain of CAMKK1 and CAMKK2 was assessed. The results are presented in **Table 1** and **Figure 5**. Generally, the ΔT_m_ values generated on CAMKK2 were higher compared to CAMKK1. This observation does not indicate that the compounds were more potent CAMKK2 inhibitors, but only that they induced a larger change to the melting temperature of the protein. In CAMKK1 five staurosporine analogues induced the largest shifts with ΔT_m_ >15 °C. Other very potent binders of CAMKK1 are the quinoline OTSSP167 and the azaindole GSK650394, which both shifted the melting temperature by >15 °C. GSK650394 induced the highest melting temperature shift in CAMKK2 (20.7 °C). The benzimidazoles crenolanib and CP-673451 and the quinoline OTSSP167 had the next highest CAMKK2 ΔT_m_ values.

**Table 1:** The inhibitors acquired and tested and their scaffold types. DSF results are a mean of n = 3 runs. IC_50_ values were generated in an 8-point full dose response assay. (* Assay interference; # n=1)

| Scaffold | Inhibitor | CAMKK1<br>$\Delta T_m$ (°C) | CAMKK2<br>$\Delta T_m$ (°C) | PoC<br>(1.0, 0.1, 0.01 μM) | | | IC<sub>50</sub><br>(nM) |
| --- | --- | --- | --- | --- | --- | --- | --- |
| 2,4-dianilinopyrimidine | AP26113 | 9.6 | 13.6 | 2 | 20 | 60 | 16 |
| 2,4-dianilinopyrimidine | ALK-IN-1 | 10.6 | 14.9 | 3 | 34 | 62 | 28 |
| 2,4-dianilinopyrimidine | TAE684 | 9.5 | 13.3 | 42 | 87 | 97 |  |
| 2,4-dianilinopyrimidine | TAE226 | 6.6 | 10.5 | 90 | 97 | 104 |  |
| 2,4-dianilinopyrimidine | CZC 25146 | 4 | 9.1 | 93 | 101 | 96 |  |
| 2-aminopyrimidine | CHIR99021 | 0.2 | 1 | 96 | 101 | 98 |  |
| 2-anilino-4-alkyl-aminopyrimidine | PF03814735 | 7 | 11 | 20 | 70 | 92 | 227 |
| 2-anilino-4-alkyl-aminopyrimidine- | BX912 | 1.7 | 7.6 | 26 | 90 | 100 |  |
| 2-anilino-4-alkyl aminopyrimidine | BX795 | 2 | 9.1 | 33 | 80 | 97 |  |
| 2-anilino-4-alkyl-aminopyrimidine | MRT67307 | 2.3 | 9.6 | 35 | 86 | 92 |  |
| 2-anilino-4-aryl-pyrimidine | AZD 3463 | 12.2 | 12.7 | 26 | 74 | 103 | 185 |
| 2-anilino-4-aryl-pyrimidine | Pacritinib | 7.4 | 10.1 | 31 | 78 | 96 |  |
| 4,6-disubstituted-pyrimidine | BGJ 398 | 0.2 | 0 | 96 | 99 | 100 |  |
| Amide hinge binder | Gilteritinib | 10 | 11.7 | 18 | 74 | 84 | 204 |
| Aminobenzothiazole (Type II) | TAK632 | 5.9 | 2.3 | 82 | 87 | 92 |  |
| Aminopyrazole | XL228 | 8.6 | 9.9 | 26 | 65 | 94 |  |
| Aminopyrazole | PF3758309 | 6.2 | 9.6 | 71 | 78 | 95 |  |
| Aminotriazole | CDK 1/2 Inhibitor III | 9.2 | 13.3 | 3 | 17 | 69 | 19 |
| Azaindole | GSK650394 | 15.8 | 20.7 | 3 | 4 | 14 | 3 |
| Benzimidazole | CP 673451 | 12.6 | 16.8 | 17 | 46 | 88 | 73 |
| Benzimidazole | Crenolanib | 13.9 | 17.6 | 47 | 92 | 118 |  |
| Benzimidazole | GSK461364 | 0 | 1.6 | 92 | 96 | 95 |  |
| Flavone | Alvocidib | 7 | 5.9 | 29 | 62 | 102 |  |
| Fused pyrimidine | BI2536 | 8.9 | 9.9 | 54 | 70 | 101 | 96 |
| Fused pyrimidine | BI6727 | 7.7 | 7.9 | 75 | 94 | 104 |  |
| Fused pyrimidine | Milciclib | 7 | 9.5 | 76 | 105 | 107 |  |
| Fused pyrimidine | MLN 0905 | 1.2 | 0.3 | 86 | 95 | 99 |  |
| Fused pyrimidine | MK1775 | 4.2 | 5.3 | 99 | 99 | 101 |  |
| Imidazopyridazine (Type II) | Ponatinib | 9.4 | 10.9 | 53 | 92 | 99 |  |
| Indazole | KW2449 | 5.3 | 6.4 | 42 | 84 | 96 |  |
| Miscellaneous | STO 609 | 7.6 | 11.3 | 21 | 56 | 63 | 58 |
| Oxindole | JAK3 Inhibitor VI | 7.8 | 12.4 | 21 | 81 | 87 | 26 |
| Oxindole | PHA665752 | * | * | 67 | 93 | 109 |  |
| Oxindole | SU6656 | * | * | 71 | 86 | 95 |  |
| Oxindole | Hesperadin | 8.6 | 11.5 | 84 | 95 | 99 |  |
| Oxindole | BIO | 13.4<sup>#</sup> | 7.2<sup>#</sup> | 89 | 100 | 102 |  |
| Purine | Aminopurvalanol | 6.9 | 8.3 | 22 | 80 | 100 |  |
| Pyrazolo-pyrimidine | LDN 193189 | 4.8 | 9.1 | 50 | 62 | 68 | 146 |
| Pyrazolo-pyrimidine | Dorsomorphin | 7.4 | 8.9 | 58 | 96 | 100 |  |
| Pyridine (Type II) | Rebasitinib | 6.4 | 8.7 | 81 | 93 | 95 |  |
| Pyrrolotriazine | Brivanib | 0.7 | 0 | 100 | 100 | 101 |  |
| Quinoline | OTSSP167 | 16.6 | 16.7 | 0 | 49 | 84 | 86 |
| Quinoline | Pelitinib | 5.8 | 8.1 | 60 | 84 | 100 |  |
| Quinolines | GSK1059615 | 2 | 6.7 | 34 | 81 | 92 |  |
| Quinolines | Ro 3306 | 0 | 5.8 | 88 | 94 | 94 |  |
| Staurosporine Analogue | Lestaurtinib | 18.4 | 16.3 | 3 | 33 | 64 | 25 |
| Staurosporine Analogue | K252a | 16.7 | 15.3 | 31 | 4 | 99 |  |
| Staurosporine Analogue | Midostaurin | 15.2 | 12.8 | 38 | 82 | 97 |  |
| Staurosporine Analogue | SB 218078 | 22.3 | 9.8 | 64 | 86 | 98 |  |
| Staurosporine Analogue | Go6976 | 9.6 | 9 | 80 | 92 | 97 |  |
| Staurosporine Analogue | Ro 31-8220 | 1.9 | 5.7 | 85 | 100 | 100 |  |
| Triazolopyridine | Filgotinib | 0 | 0.5 | 72 | 86 | 96 |  |

**Figure 5:**
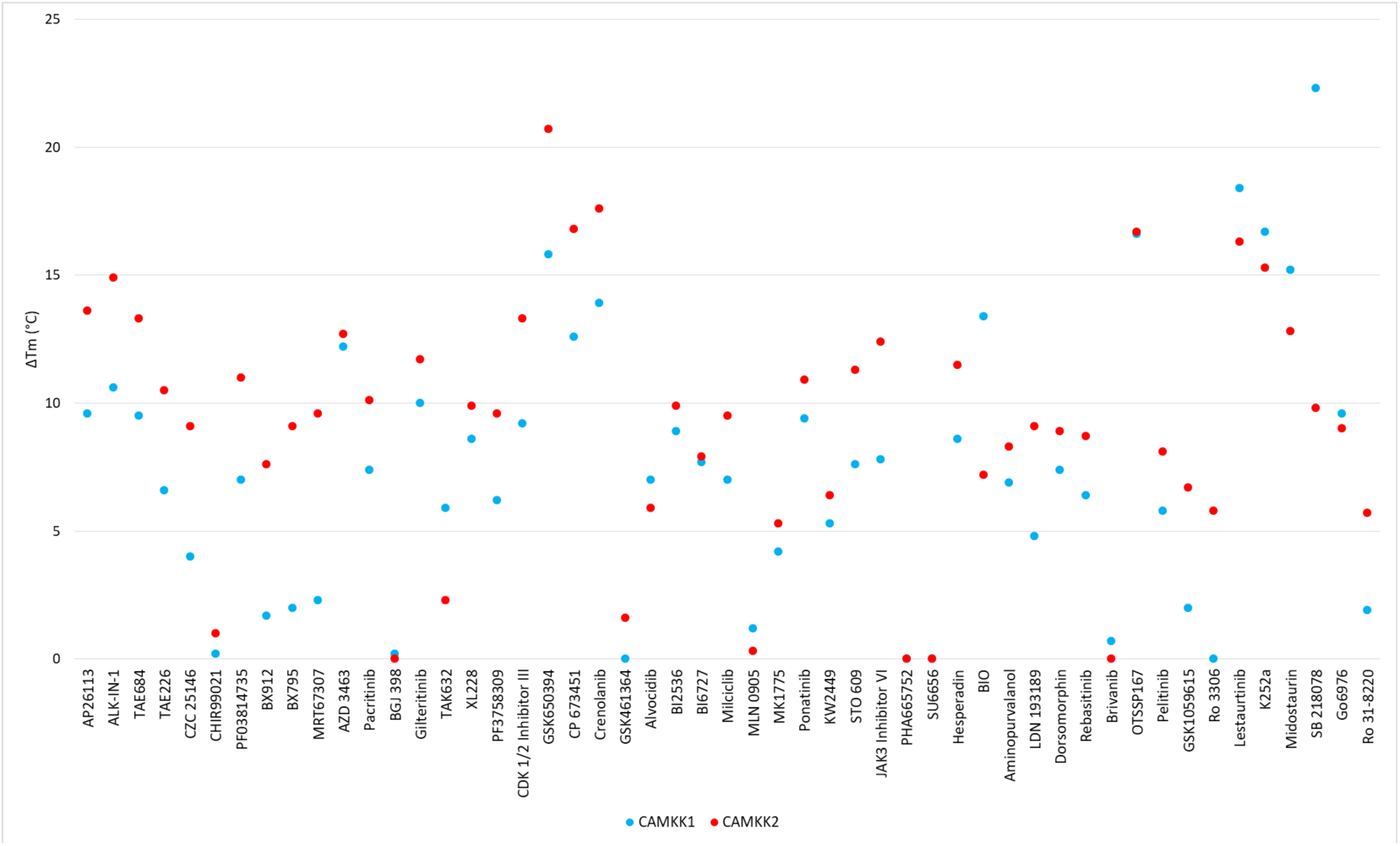
DSF results for literature inhibitors tested on CAMKK1 and CAMKK2. Results are the mean of 3 repeats run simultaneously under identical conditions. Compounds that yielded negative values are presented as having ΔT_m_ = 0 °C.

Compounds were tested in a CAMKK2 enzyme inhibition assay at 0.01, 0.1 and 1 µM concentrations (**Table 1**). STO-609 resulted 21% activity remaining (PoC) at 1 μM and generated an IC_50_ of 56 nM when tested in a full 8-point dose response assay. At 1 μM ten compounds had lower PoC values than STO-609, representing a range of different chemotypes. IC_50_ values were generated for these compounds, and also for AZD3463, crenolanib and BI2536, three inhibitors which had been co-crystallized with CAMKK2.[130] The ten most potent CAMKK2 inhibitors are shown in **Figure 6**. Their IC_50_ values correlated well with published values where available.[131]

**Figure 6:**
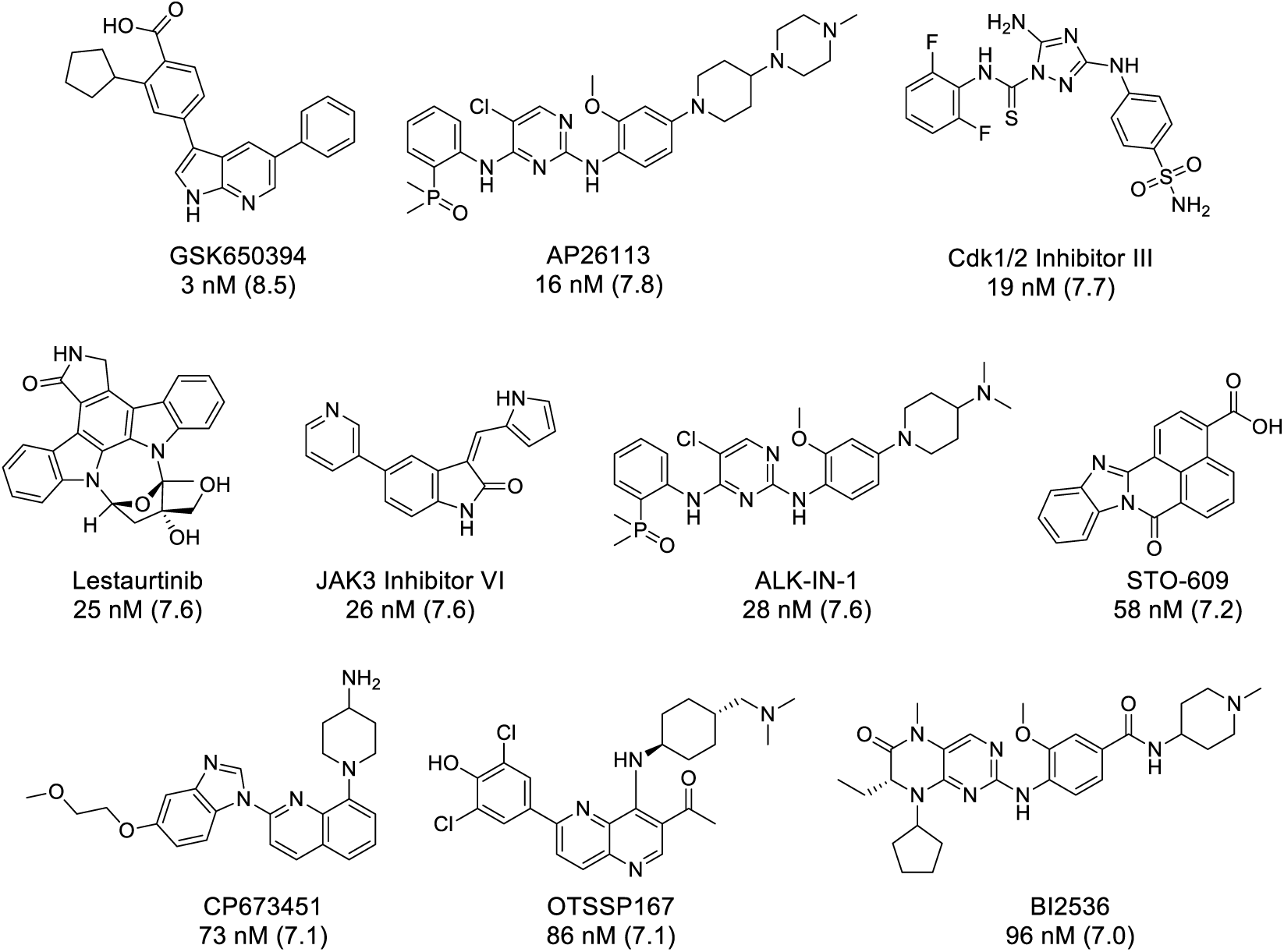
The 10 most potent CAMKK2 inhibitors identified in our screen. pIC50 values are shown in parentheses.

The IC_50_ for GSK650394 was 3 nM (pIC_50_ = 8.5), in line with the activity reported in the literature (pIC_50_= 9.2) and was the most potent CAMKK2 inhibitor identified in this study. Kinase selectivity was calculated using publicly available data and these data are presented in **Table 2**. For data from single concentration screening the number of kinases inhibited by greater than 90% (or PoC <10%) at the given screening concentration was divided by the number of kinases in the screening panel to generate an S_10_ selectivity value. If IC_50_ data was available, the selectivity metric was calculated by dividing the number of kinases with an IC_50_ below the chosen threshold by the number of kinases screened. Caution should be used in comparing these selectivity values, since data was collected using different assay formats, various inhibitor concentrations, and reported in different formats (for example PoC, K_d_ values, and IC_50_ values). However, the results in **Table 2** do provide a useful qualitative assessment of kinase selectivity.

**Table 2:** Potent CAMKK2 inhibitors, their IC_50_ values and calculated selectivity metrics. The method for determination of a given metric was calculated is shown under the method column. The number of kinases used in the calculation is shown in parentheses. The origin of the data is shown under source.

| Inhibitor | IC<sub>50</sub> (nM) | Selectivity metric | Method (#kinases in panel) | Source |
| --- | --- | --- | --- | --- |
| GSK650394 | 3 | 0.083 | S<sub>10</sub> @ 1 μM (334) | RBC |
| AP26113 | 16 | 0.125 | IC<sub>50</sub> ≤50 nM (289) | Paper[132] |
| CDK 1/2 Inhibitor III | 19 | 0.278 | S<sub>10</sub> @ 0.5 μM (300) | RBC |
| Lestaurtinib | 25 | 0.356 | K<sub>d</sub> ≤50 nM (443) | LINCS |
| JAK3 Inhibitor VI | 26 | 0.163 | S<sub>10</sub> @ 0.5 μM (300) | RBC |
| ALK-IN-1 | 28 | - | - | - |
| STO 609 | 58 | 0.015 | S<sub>10</sub> @ 1 μM (409) | KINOMEscan |
| CP 673451 | 73 | - | - | - |
| OTSSP167 | 86 | 0.66 | S<sub>10</sub> @ 1 μM (141) | MRC |
| BI2536 | 96 | 0.007 | S<sub>10</sub> @ 1 μM (131) | MRC |

## 3. Discussion

When profiled at MRC, at 1 µM GSK650394 inhibited 7 of 85 kinases by more than 90%, equating to an S_10_ value of 0.082. In a larger a panel of 334 kinases (Reaction Biology Corporation), GSK650394 inhibited 29 kinases by more than 90%, representing a nearly identical S_10_ value of 0.083 (**Table S3**). The number of kinase off-targets limits the utility of GSK650394 as a chemical probe for CAMKK2, since attributing a phenotypic response to a single kinase would be precarious. The S_10_ metric could be calculated for STO-609, BI2536 and OTSSP167 at 1 µM, and for CDK 1/2 inhibitor III and JAK3 inhibitor VI, at 0.5 µM. STO-609 had moderate selectivity with an S_10_ at 1uM of 0.015. In this set, BI2536 appears to be the most selective of the CAMKK2 inhibitors. In the literature BI2536 was reported as a potent PLK1, PLK2 and PLK3 inhibitor, with Kd’s <10 nM.[133]. In addition to the collateral PLK inhibition, BI2536 has been shown to be a potent inhibitor of BRD4, which is a critical regulator of cell division and confounds its use as a selective kinase inhibitor probe.[134] OTSSP167 is an extremely potent but promiscuous kinase inhibitor, with an S_10_ at 1 μM of 0.66. Neither CDK1/2 inhibitor III or JAK3 inhibitor VI can be considered selective, even at a lower concentration of 0.5 µM, they have S_10_ values of 0.278 and 0.163 respectively. The S_10_ metric could not be calculated for AP26113, but IC_50_ values were generated against a panel of 289 kinases.[135] From this panel, AP26113 generated IC_50_ values below 10 nM for 10 kinases, and below 50 nM for 24 kinases. Thus, it is likely to potently inhibit almost 10% of the kinome. Kinases inhibited by AP26113 included known oncology targets such as FER, EFGR and FLT3 and the polypharmacology of this compound may contribute to its efficacy in patients. ALK-IN-1 is an analogue of AP26113, differing only in the solvent exposed moiety. Although kinome profiling data was not available for ALK-IN-1, it may show polypharmacology similar to AP26113. Lestaurtinib was profiled in the KINOME*scan* panel and was extremely promiscuous, demonstrating K_d_’s <20 nM for 111 out of 443 kinases. There was no profiling data available for CP673451 but the S_10_ value for crenolanib, a close analogue, was 0.142.

The data in **Table 2** revealed several potential starting points for development of potent and selective CAMKK2 inhibitors. GSK650394 is potent but suffers from poor kinase selectivity. Several strategies could be utilized to increase the compound’s kinase selectivity. Price *et al* demonstrated that replacement of the carboxylic acid group by an amide retained CAMKK2 potency.[89] The authors also demonstrate that the cyclopentyl group occupied a pocket that tolerated alternative substitutions, although no group larger than the cyclopentyl was reported. The unsubstituted phenyl ring could also be functionalized, and there is likely room to grow this into the solvent exposed region where kinases tend to be less highly conserved. A promising strategy would be bioisosteric replacement of the azaindole core, since it is known that the hydrogen bond interactions between a kinase inhibitor and the enzyme’s hinge region are important for selectivity and potency.[136]

BI2536 represents another potential starting point to develop a potent and selective CAMKK2 inhibitor as the number of off targets is limited. The activity of the compound on the PLK family could be addressed by modifying the core, specifically by further functionalizing the fused piperazinone ring. The *N*-methyl could be replaced with different alkyl groups, which may also be applied to the ethyl group. There is room to extend the compound at the ethyl and potentially install a group that can interact with the lysine that forms a H-bond with the acid moiety in GSK650394. The cyclopentyl group of BI2536 also occupies the same pocket as identified in GSK650394 and would be an interesting group to modify.[131] Modifications of these groups could induce selectivity for CAMKK2 over the PLKs. However, it will also be important to remove all the off-target bromodomain activity of this chemotype. Although we identified other potent scaffolds, their promiscuity is severe, and they do not represent good starting points for the development of selective inhibitors.

## Conclusion

CAMKK2 is an extremely important kinase in cell signaling, playing an essential role in key cellular processes. Deregulation of CAMKK2 has implications in oncological, metabolic and neurological diseases. There is currently no high quality CAMKK2 chemical probe available to the scientific community for use in target validation and pre-clinical translation studies. The need for a selective CAMKK2 inhibitor is acute, given that the most widely studied inhibitor, STO-609, has poor solubility and shows collateral inhibition of several other regulatory kinases. Through this review of the literature and public databases and generation of new screening data, additional starting points for the development of selective CAMKK2 inhibitors have been identified and strategies to optimize them have been proposed. We anticipate that these results will help the community make further progress towards CAMKK2 inhibitors that will delineate roles played by CAMKK2 in disease and can be used to explore its inhibition in a clinical setting.

## 4. Materials and Methods

### CAMKK1 and CAMKK2 DSF assay

Small molecule screening by DSF were performed as described previously.[137, 138] Briefly the DSF assay was performed in 96-well format. Purified CAMKK1 or CAMKK2 was diluted to 2 µM kinase in 100 mM potassium phosphate pH 7.5, 150 mM NaCl, and 10% glycerol supplemented with 5x SYPRO Orange (Invitrogen). All assay experiments used 19.5 µL of 2 µM kinase and SYPRO Orange mixture. Compounds solubilized in DMSO were used at 12.5 µM final concentration, with a 2.5% concentration of DMSO per well. PCR plates were sealed using optically clear films and transferred to a C1000 thermal cycler with CFX-96 RT-PCR head (BioRad). The fluorescence intensity was measured over a temperature gradient from 25– 95 °C at a constant rate of 0.05 °C/s. Curve fitting and protein melting temperatures were calculated based on a Boltzmann function fitting to experimental data (GraphPad Prism 8). Protein with the addition of 2.5% DMSO was used as a reference. All experiments were carried out in triplicate and the mean of the ΔT_m_ is reported. Compounds that provided negative values are presented as having a ΔT_m_ of 0 °C.

### CAMKK2 enzyme assay

CAMKK2 activity was determined by measuring the transfer of radiolabeled phosphate from [γ-^32^P]-ATP to a synthetic peptide substrate (CaMKKtide) as previously described.(38) Briefly, purified recombinant CAMKK2 (100 pM) was incubated in assay buffer (50 mM HEPES [pH 7.4], 1 mM DTT, 0.02% [v/v] Brij-35) containing 200 µM CaMKKtide (Genscript), 100 µM CaCl_2_, 1 µM CaM (Sigma-Aldrich), 200 µM [γ-^32^P]-ATP (Perkin Elmer), 5 mM MgCl_2_ (Sigma-Aldrich) and various concentrations of inhibitors (0-1 µM) in a standard 30 µl assay for 10 min at 30 °C. Reactions were terminated by spotting 15 µl onto P81 phosphocellulose paper (GE Lifesciences) and washing extensively in 1% phosphoric acid (Sigma-Aldrich). Radioactivity was quantified by liquid scintillation counting.

## Supplementary Materials

The following are available online, **Table S1**: KINOME*scan* data for STO-609; **Table S2**: Names and SMILES strings of compounds tested in this manuscript; **Table S3**: RBC data for GSK650394;

## Author Contributions

SNOB wrote the manuscript; DHD & SNOB conception & design; JWS performed enzymology testing; JRP performed the DSF experiments; CGL expressed and purified CaMKK2 for enzymology testing; ASS performed protein expression & purification for the DSF; SNOB, DHD, CIW database mining; SNOB, DHD & CIW – identified literature hits; RMC contributed to structural analysis; SNOB, DHD, CIW, JWS, JSO, TMW & BJE edited the manuscript; DHD – project administration; DHD – funding acquisition.

## References

1. Chin, D.; Means, A. R., Calmodulin: a prototypical calcium sensor. Trends Cell Biol 2000, 10, (8), 322–8.

2. Hook, S. S.; Means, A. R., Ca(2+)/CaM-dependent kinases: from activation to function. Annu Rev Pharmacol Toxicol 2001, 41, (1), 471–505.

3. Chin, D.; Means, A. R., Calmodulin: a prototypical calcium sensor. Trends in Cell Biology 2000, 10, (8), 322–328.

4. Sharma, R. K.; Parameswaran, S., Calmodulin-binding proteins: A journey of 40 years. Cell Calcium 2018, 75, 89–100.

5. Shen, X.; Valencia, C. A.; Szostak, J. W.; Dong, B.; Liu, R., Scanning the human proteome for calmodulin-binding proteins. Proc Natl Acad Sci U S A 2005, 102, (17), 5969–74.

6. Tidow, H.; Nissen, P., Structural diversity of calmodulin binding to its target sites. FEBS J 2013, 280, (21), 5551–65.

7. Hanks, S. K.; Hunter, T., Protein kinases 6. The eukaryotic protein kinase superfamily: kinase (catalytic) domain structure and classification. FASEB J 1995, 9, (8), 576–96.

8. Manning, G.; Whyte, D. B.; Martinez, R.; Hunter, T.; Sudarsanam, S., The protein kinase complement of the human genome. Science 2002, 298, (5600), 1912–34.

9. Bayer, K. U.; Schulman, H., CaM Kinase: Still Inspiring at 40. Neuron 2019, 103, (3), 380–394.

10. Soderling, T. R., The Ca2+-calmodulin-dependent protein kinase cascade. Trends in Biochemical Sciences 1999, 24, (6), 232–236.

11. de Souza Almeida Matos, A. L.; Oakhill, J. S.; Moreira, J.; Loh, K.; Galic, S.; Scott, J. W., Allosteric regulation of AMP-activated protein kinase by adenylate nucleotides and small-molecule drugs. Biochem Soc Trans 2019, 47, (2), 733–741.

12. Racioppi, L.; Means, A. R., Calcium/calmodulin-dependent protein kinase kinase 2: roles in signaling and pathophysiology. J Biol Chem 2012, 287, (38), 31658–65.

13. Marcelo, K. L.; Means, A. R.; York, B., The Ca(2+)/Calmodulin/CaMKK2 Axis: Nature’s Metabolic CaMshaft. Trends Endocrinol Metab 2016, 27, (10), 706–718.

14. Anderson, K. A.; Means, R. L.; Huang, Q. H.; Kemp, B. E.; Goldstein, E. G.; Selbert, M. A.; Edelman, A. M.; Fremeau, R. T.; Means, A. R., Components of a calmodulin-dependent protein kinase cascade. Molecular cloning, functional characterization and cellular localization of Ca2+/calmodulin-dependent protein kinase kinase beta. J Biol Chem 1998, 273, (48), 31880–9.

15. Hawley, S. A.; Selbert, M. A.; Goldstein, E. G.; Edelman, A. M.; Carling, D.; Hardie, D. G., 5’-AMP activates the AMP-activated protein kinase cascade, and Ca2+/calmodulin activates the calmodulin-dependent protein kinase I cascade, via three independent mechanisms. J Biol Chem 1995, 270, (45), 27186–91.

16. Yano, S.; Tokumitsu, H.; Soderling, T. R., Calcium promotes cell survival through CaM-K kinase activation of the protein-kinase-B pathway. Nature 1998, 396, (6711), 584–7.

17. Gocher, A. M.; Azabdaftari, G.; Euscher, L. M.; Dai, S.; Karacosta, L. G.; Franke, T. F.; Edelman, A. M., Akt activation by Ca(2+)/calmodulin-dependent protein kinase kinase 2 (CaMKK2) in ovarian cancer cells. J Biol Chem 2017, 292, (34), 14188–14204.

18. Marcelo, K. L.; Means, A. R.; York, B., The Ca(2+)/Calmodulin/CaMKK2 Axis: Nature’s Metabolic CaMshaft. Trends Endocrinol Metab 2016, 27, (10), 706–718.

19. Bellacosa, A.; Testa, J. R.; Moore, R.; Larue, L., A portrait of AKT kinases: human cancer and animal models depict a family with strong individualities. Cancer Biol Ther 2004, 3, (3), 268–75.

20. Racioppi, L.; Means, A. R., Calcium/Calmodulin-dependent Protein Kinase Kinase 2: Roles in Signaling and Pathophysiology. 2012, 287, (38), 31658–31665.

21. Picciotto, M. R.; Zoli, M.; Bertuzzi, G.; Nairn, A. C., Immunochemical localization of calcium/calmodulin-dependent protein kinase I. Synapse 1995, 20, (1), 75–84.

22. Kamata, A.; Sakagami, H.; Tokumitsu, H.; Owada, Y.; Fukunaga, K.; Kondo, H., Spatiotemporal expression of four isoforms of Ca2+/calmodulin-dependent protein kinase I in brain and its possible roles in hippocampal dendritic growth. Neurosci Res 2007, 57, (1), 86–97.

23. Joseph, J. D.; Means, A. R., Identification and characterization of two Ca2+/CaM-dependent protein kinases required for normal nuclear division in Aspergillus nidulans. J Biol Chem 2000, 275, (49), 38230–8.

24. Skelding, K. A.; Rostas, J. A.; Verrills, N. M., Controlling the cell cycle: the role of calcium/calmodulin-stimulated protein kinases I and II. Cell Cycle 2011, 10, (4), 631–9.

25. Wayman, G. A.; Kaech, S.; Grant, W. F.; Davare, M.; Impey, S.; Tokumitsu, H.; Nozaki, N.; Banker, G.; Soderling, T. R., Regulation of axonal extension and growth cone motility by calmodulin-dependent protein kinase I. J Neurosci 2004, 24, (15), 3786–94.

26. Ang, E. S.; Zhang, P.; Steer, J. H.; Tan, J. W.; Yip, K.; Zheng, M. H.; Joyce, D. A.; Xu, J., Calcium/calmodulin-dependent kinase activity is required for efficient induction of osteoclast differentiation and bone resorption by receptor activator of nuclear factor kappa B ligand (RANKL). J Cell Physiol 2007, 212, (3), 787–95.

27. Condon, J. C.; Pezzi, V.; Drummond, B. M.; Yin, S.; Rainey, W. E., Calmodulin-dependent kinase I regulates adrenal cell expression of aldosterone synthase. Endocrinology 2002, 143, (9), 3651–7.

28. Schmitt, J. M.; Guire, E. S.; Saneyoshi, T.; Soderling, T. R., Calmodulin-dependent kinase kinase/calmodulin kinase I activity gates extracellular-regulated kinase-dependent long-term potentiation. J Neurosci 2005, 25, (5), 1281–90.

29. Takemoto-Kimura, S.; Ageta-Ishihara, N.; Nonaka, M.; Adachi-Morishima, A.; Mano, T.; Okamura, M.; Fujii, H.; Fuse, T.; Hoshino, M.; Suzuki, S.; Kojima, M.; Mishina, M.; Okuno, H.; Bito, H., Regulation of dendritogenesis via a lipid-raft-associated Ca2+/calmodulin-dependent protein kinase CLICK-III/CaMKIgamma. Neuron 2007, 54, (5), 755–70.

30. Ohmstede, C. A.; Jensen, K. F.; Sahyoun, N. E., Ca2+/calmodulin-dependent protein kinase enriched in cerebellar granule cells. Identification of a novel neuronal calmodulin-dependent protein kinase. J Biol Chem 1989, 264, (10), 5866–75.

31. Kitsos, C. M.; Sankar, U.; Illario, M.; Colomer-Font, J. M.; Duncan, A. W.; Ribar, T. J.; Reya, T.; Means, A. R., Calmodulin-dependent protein kinase IV regulates hematopoietic stem cell maintenance. J Biol Chem 2005, 280, (39), 33101–8.

32. Wu, J. Y.; Means, A. R., Ca(2+)/calmodulin-dependent protein kinase IV is expressed in spermatids and targeted to chromatin and the nuclear matrix. J Biol Chem 2000, 275, (11), 7994–9.

33. Wu, J. Y.; Gonzalez-Robayna, I. J.; Richards, J. S.; Means, A. R., Female fertility is reduced in mice lacking Ca2+/calmodulin-dependent protein kinase IV. Endocrinology 2000, 141, (12), 4777–83.

34. Skelding, K. A.; Rostas, J. A. P.; Verrills, N. M., Controlling the cell cycle: The role of calcium/calmodulin-stimulated protein kinases I and II. Cell Cycle 2011, 10, (4), 631–639.

35. Kimura, Y.; Corcoran, E. E.; Eto, K.; Gengyo-Ando, K.; Muramatsu, M. A.; Kobayashi, R.; Freedman, J. H.; Mitani, S.; Hagiwara, M.; Means, A. R.; Tokumitsu, H., A CaMK cascade activates CRE-mediated transcription in neurons of Caenorhabditis elegans. EMBO Rep 2002, 3, (10), 962–6.

36. Bleier, J.; Toliver, A., Exploring the Role of CaMKIV in Homeostatic Plasticity. J Neurosci 2017, 37, (48), 11520–11522.

37. Takemura, M.; Mishima, T.; Wang, Y.; Kasahara, J.; Fukunaga, K.; Ohashi, K.; Mizuno, K., Ca2+/calmodulin-dependent protein kinase IV-mediated LIM kinase activation is critical for calcium signal-induced neurite outgrowth. J Biol Chem 2009, 284, (42), 28554–62.

38. Wei, F.; Qiu, C. S.; Liauw, J.; Robinson, D. A.; Ho, N.; Chatila, T.; Zhuo, M., Calcium calmodulin-dependent protein kinase IV is required for fear memory. Nat Neurosci 2002, 5, (6), 573–9.

39. Racioppi, L.; Means, A. R., Calcium/calmodulin-dependent kinase IV in immune and inflammatory responses: novel routes for an ancient traveller. Trends Immunol 2008, 29, (12), 600–7.

40. Hardie, D. G., AMP-activated/SNF1 protein kinases: conserved guardians of cellular energy. Nat Rev Mol Cell Biol 2007, 8, (10), 774–85.

41. Mihaylova, M. M.; Shaw, R. J., The AMPK signalling pathway coordinates cell growth, autophagy and metabolism. Nat Cell Biol 2011, 13, (9), 1016–23.

42. Herzig, S.; Shaw, R. J., AMPK: guardian of metabolism and mitochondrial homeostasis. Nat Rev Mol Cell Biol 2018, 19, (2), 121–135.

43. Bellacosa, A.; Testa, J. R.; moore, r.; Larue, L., A Portrait of AKT Kinases: Human Cancer and Animal Models Depict a Family with Strong Individualities. Cancer Biology & Therapy 2004, 3, (3), 268–275.

44. Nitulescu, G. M.; Van De Venter, M.; Nitulescu, G.; Ungurianu, A.; Juzenas, P.; Peng, Q.; Olaru, O. T.; Gradinaru, D.; Tsatsakis, A.; Tsoukalas, D.; Spandidos, D. A.; Margina, D., The Akt pathway in oncology therapy and beyond (Review). Int J Oncol 2018, 53, (6), 2319–2331.

45. Manning, B. D.; Toker, A., AKT/PKB Signaling: Navigating the Network. Cell 2017, 169, (3), 381–405.

46. Song, M.; Bode, A. M.; Dong, Z.; Lee, M. H., AKT as a Therapeutic Target for Cancer. Cancer Res 2019, 79, (6), 1019–1031.

47. Ishikawa, Y.; Tokumitsu, H.; Inuzuka, H.; Murata-Hori, M.; Hosoya, H.; Kobayashi, R., Identification and characterization of novel components of a Ca2+/calmodulin-dependent protein kinase cascade in HeLa cells. FEBS Lett 2003, 550, (1-3), 57-63.

48. Hsu, L. S.; Chen, G. D.; Lee, L. S.; Chi, C. W.; Cheng, J. F.; Chen, J. Y., Human Ca2+/calmodulin-dependent protein kinase kinase beta gene encodes multiple isoforms that display distinct kinase activity. J Biol Chem 2001, 276, (33), 31113–23.

49. Santiago, A. D. S.; Counago, R. M.; Ramos, P. Z.; Godoi, P. H. C.; Massirer, K. B.; Gileadi, O.; Elkins, J. M., Structural Analysis of Inhibitor Binding to CAMKK1 Identifies Features Necessary for Design of Specific Inhibitors. Sci Rep 2018, 8, (1), 14800.

50. Tokumitsu, H.; Iwabu, M.; Ishikawa, Y.; Kobayashi, R., Differential regulatory mechanism of Ca2+/calmodulin-dependent protein kinase kinase isoforms. Biochemistry 2001, 40, (46), 13925–32.

51. Edelman, A. M.; Mitchelhill, K. I.; Selbert, M. A.; Anderson, K. A.; Hook, S. S.; Stapleton, D.; Goldstein, E. G.; Means, A. R.; Kemp, B. E., Multiple Ca(2+)-calmodulin-dependent protein kinase kinases from rat brain. Purification, regulation by Ca(2+)-calmodulin, and partial amino acid sequence. J Biol Chem 1996, 271, (18), 10806–10.

52. Anderson, K. A.; Means, R. L.; Huang, Q.-H.; Kemp, B. E.; Goldstein, E. G.; Selbert, M. A.; Edelman, A. M.; Fremeau, R. T.; Means, A. R., Components of a Calmodulin-dependent Protein Kinase Cascade: MOLECULAR CLONING, FUNCTIONAL CHARACTERIZATION AND CELLULAR LOCALIZATION OF Ca2+/CALMODULIN-DEPENDENT PROTEIN KINASE KINASE β. 1998, 273, (48), 31880–31889.

53. Green, M. F.; Scott, J. W.; Steel, R.; Oakhill, J. S.; Kemp, B. E.; Means, A. R., Ca2+/Calmodulin-dependent protein kinase kinase beta is regulated by multisite phosphorylation. J Biol Chem 2011, 286, (32), 28066–79.

54. Scott, J. W.; Park, E.; Rodriguiz, R. M.; Oakhill, J. S.; Issa, S. M. A.; O’Brien, M. T.; Dite, T. A.; Langendorf, C. G.; Wetsel, W. C.; Means, A. R.; Kemp, B. E., Autophosphorylation of CaMKK2 generates autonomous activity that is disrupted by a T85S mutation linked to anxiety and bipolar disorder. Scientific Reports 2015, 5, 14436.

55. Scott, J. W.; Park, E.; Rodriguiz, R. M.; Oakhill, J. S.; Issa, S. M. A.; O’Brien, M. T.; Dite, T. A.; Langendorf, C. G.; Wetsel, W. C.; Means, A. R.; Kemp, B. E., Autophosphorylation of CaMKK2 generates autonomous activity that is disrupted by a T85S mutation linked to anxiety and bipolar disorder. Scientific Reports 2015, 5, 14436.

56. O’Brien, M. T.; Oakhill, J. S.; Ling, N. X.; Langendorf, C. G.; Hoque, A.; Dite, T. A.; Means, A. R.; Kemp, B. E.; Scott, J. W., Impact of Genetic Variation on Human CaMKK2 Regulation by Ca(2+)-Calmodulin and Multisite Phosphorylation. Sci Rep 2017, 7, 43264.

57. Wayman, G. A.; Tokumitsu, H.; Soderling, T. R., Inhibitory cross-talk by cAMP kinase on the calmodulin-dependent protein kinase cascade. J Biol Chem 1997, 272, (26), 16073–6.

58. Santiago, A. d. S.; Couñago, R. M.; Ramos, P. Z.; Godoi, P. H. C.; Massirer, K. B.; Gileadi, O.; Elkins, J. M., Structural Analysis of Inhibitor Binding to CAMKK1 Identifies Features Necessary for Design of Specific Inhibitors. Scientific Reports 2018, 8, (1), 14800.

59. Price, D. J.; Drewry, D. H.; Schaller, L. T.; Thompson, B. D.; Reid, P. R.; Maloney, P. R.; Liang, X.; Banker, P.; Buckholz, R. G.; Selley, P. K.; McDonald, O. B.; Smith, J. L.; Shearer, T. W.; Cox, R. F.; Williams, S. P.; Reid, R. A.; Tacconi, S.; Faggioni, F.; Piubelli, C.; Sartori, I.; Tessari, M.; Wang, T. Y., An orally available, brain-penetrant CAMKK2 inhibitor reduces food intake in rodent model. Bioorg Med Chem Lett 2018, 28, (10), 1958–1963.

60. Kukimoto-Niino, M.; Yoshikawa, S.; Takagi, T.; Ohsawa, N.; Tomabechi, Y.; Terada, T.; Shirouzu, M.; Suzuki, A.; Lee, S.; Yamauchi, T.; Okada-Iwabu, M.; Iwabu, M.; Kadowaki, T.; Minokoshi, Y.; Yokoyama, S., Crystal structure of the Ca(2)(+)/calmodulin-dependent protein kinase kinase in complex with the inhibitor STO-609. J Biol Chem 2011, 286, (25), 22570–9.

61. Asquith, C. R. M.; Godoi, P. H.; Counago, R. M.; Laitinen, T.; Scott, J. W.; Langendorf, C. G.; Oakhill, J. S.; Drewry, D. H.; Zuercher, W. J.; Koutentis, P. A.; Willson, T. M.; Kalogirou, A. S., 1,2,6-Thiadiazinones as Novel Narrow Spectrum Calcium/Calmodulin-Dependent Protein Kinase Kinase 2 (CaMKK2) Inhibitors. Molecules 2018, 23, (5).

62. Massie, C. E.; Lynch, A.; Ramos-Montoya, A.; Boren, J.; Stark, R.; Fazli, L.; Warren, A.; Scott, H.; Madhu, B.; Sharma, N.; Bon, H.; Zecchini, V.; Smith, D. M.; Denicola, G. M.; Mathews, N.; Osborne, M.; Hadfield, J.; Macarthur, S.; Adryan, B.; Lyons, S. K.; Brindle, K. M.; Griffiths, J.; Gleave, M. E.; Rennie, P. S.; Neal, D. E.; Mills, I. G., The androgen receptor fuels prostate cancer by regulating central metabolism and biosynthesis. EMBO J 2011, 30, (13), 2719–33.

63. Karacosta, L. G.; Foster, B. A.; Azabdaftari, G.; Feliciano, D. M.; Edelman, A. M., A regulatory feedback loop between Ca2+/calmodulin-dependent protein kinase kinase 2 (CaMKK2) and the androgen receptor in prostate cancer progression. J Biol Chem 2012, 287, (29), 24832–43.

64. Shima, T.; Mizokami, A.; Miyagi, T.; Kawai, K.; Izumi, K.; Kumaki, M.; Ofude, M.; Zhang, J.; Keller, E. T.; Namiki, M., Down-regulation of calcium/calmodulin-dependent protein kinase kinase 2 by androgen deprivation induces castration-resistant prostate cancer. Prostate 2012, 72, (16), 1789–801.

65. Subbannayya, Y.; Syed, N.; Barbhuiya, M. A.; Raja, R.; Marimuthu, A.; Sahasrabuddhe, N.; Pinto, S. M.; Manda, S. S.; Renuse, S.; Manju, H. C.; Zameer, M. A.; Sharma, J.; Brait, M.; Srikumar, K.; Roa, J. C.; Vijaya Kumar, M.; Kumar, K. V.; Prasad, T. S.; Ramaswamy, G.; Kumar, R. V.; Pandey, A.; Gowda, H.; Chatterjee, A., Calcium calmodulin dependent kinase kinase 2 - a novel therapeutic target for gastric adenocarcinoma. Cancer Biol Ther 2015, 16, (2), 336–45.

66. Lin, F.; Marcelo, K. L.; Rajapakshe, K.; Coarfa, C.; Dean, A.; Wilganowski, N.; Robinson, H.; Sevick, E.; Bissig, K. D.; Goldie, L. C.; Means, A. R.; York, B., The camKK2/camKIV relay is an essential regulator of hepatic cancer. Hepatology 2015, 62, (2), 505–20.

67. Liu, D. M.; Wang, H. J.; Han, B.; Meng, X. Q.; Chen, M. H.; Yang, D. B.; Sun, Y.; Li, Y. L.; Jiang, C. L., CAMKK2, Regulated by Promoter Methylation, is a Prognostic Marker in Diffuse Gliomas. CNS Neurosci Ther 2016, 22, (6), 518–24.

68. Gocher, A. M.; Azabdaftari, G.; Euscher, L. M.; Dai, S.; Karacosta, L. G.; Franke, T. F.; Edelman, A. M., Akt activation by Ca2+/calmodulin-dependent protein kinase kinase 2 (CaMKK2) in ovarian cancer cells. 2017, 292, (34), 14188–14204.

69. Rodriguez-Mora, O. G.; LaHair, M. M.; McCubrey, J. A.; Franklin, R. A., Calcium/calmodulin-dependent kinase I and calcium/calmodulin-dependent kinase kinase participate in the control of cell cycle progression in MCF-7 human breast cancer cells. Cancer Res 2005, 65, (12), 5408–16.

70. Schmitt, J. M.; Abell, E.; Wagner, A.; Davare, M. A., ERK activation and cell growth require CaM kinases in MCF-7 breast cancer cells. Mol Cell Biochem 2010, 335, (1-2), 155-71.

71. Karacosta, L. G.; Foster, B. A.; Azabdaftari, G.; Feliciano, D. M.; Edelman, A. M., A Regulatory Feedback Loop Between Ca2+/Calmodulin-dependent Protein Kinase Kinase 2 (CaMKK2) and the Androgen Receptor in Prostate Cancer Progression. 2012, 287, (29), 24832–24843.

72. Davare, M. A.; Saneyoshi, T.; Soderling, T. R., Calmodulin-kinases regulate basal and estrogen stimulated medulloblastoma migration via Rac1. J Neurooncol 2011, 104, (1), 65–82.

73. Frigo, D. E.; Howe, M. K.; Wittmann, B. M.; Brunner, A. M.; Cushman, I.; Wang, Q. B.; Brown, M.; Means, A. R.; McDonnell, D. P., CaM Kinase Kinase beta-Mediated Activation of the Growth Regulatory Kinase AMPK Is Required for Androgen-Dependent Migration of Prostate Cancer Cells. Cancer Research 2011, 71, (2), 528–537.

74. Fu, H.; He, H. C.; Han, Z. D.; Wan, Y. P.; Luo, H. W.; Huang, Y. Q.; Cai, C.; Liang, Y. X.; Dai, Q. S.; Jiang, F. N.; Zhong, W. D., MicroRNA-224 and its target CAMKK2 synergistically influence tumor progression and patient prognosis in prostate cancer. Tumour Biol 2015, 36, (3), 1983–91.

75. Lin, F.; Marcelo, K. L.; Rajapakshe, K.; Coarfa, C.; Dean, A.; Wilganowski, N.; Robinson, H.; Sevick, E.; Bissig, K.-D.; Goldie, L. C.; Means, A. R.; York, B., The camKK2/camKIV relay is an essential regulator of hepatic cancer. 2015, 62, (2), 505–520.

76. Liu, D.-M.; Wang, H.-J.; Han, B.; Meng, X.-Q.; Chen, M.-H.; Yang, D.-B.; Sun, Y.; Li, Y.-L.; Jiang, C.-L., CAMKK2, Regulated by Promoter Methylation, is a Prognostic Marker in Diffuse Gliomas. 2016, 22, (6), 518–524.

77. Ma, Z.; Wen, D.; Wang, X.; Yang, L.; Liu, T.; Liu, J.; Zhu, J.; Fang, X., Growth inhibition of human gastric adenocarcinoma cells in vitro by STO-609 is independent of calcium/calmodulin-dependent protein kinase kinase-beta and adenosine monophosphate-activated protein kinase. Am J Transl Res 2016, 8, (2), 1164–71.

78. Massie, C. E.; Lynch, A.; Ramos-Montoya, A.; Boren, J.; Stark, R.; Fazli, L.; Warren, A.; Scott, H.; Madhu, B.; Sharma, N.; Bon, H.; Zecchini, V.; Smith, D. M.; DeNicola, G. M.; Mathews, N.; Osborne, M.; Hadfield, J.; MacArthur, S.; Adryan, B.; Lyons, S. K.; Brindle, K. M.; Griffiths, J.; Gleave, M. E.; Rennie, P. S.; Neal, D. E.; Mills, I. G., The androgen receptor fuels prostate cancer by regulating central metabolism and biosynthesis. 2011, 30, (13), 2719–2733.

79. Rodriguez-Mora, O. G.; LaHair, M. M.; McCubrey, J. A.; Franklin, R. A., Calcium/Calmodulin-Dependent Kinase I and Calcium/Calmodulin-Dependent Kinase Kinase Participate in the Control of Cell Cycle Progression in MCF-7 Human Breast Cancer Cells. 2005, 65, (12), 5408–5416.

80. Shima, T.; Mizokami, A.; Miyagi, T.; Kawai, K.; Izumi, K.; Kumaki, M.; Ofude, M.; Zhang, J.; Keller, E. T.; Namiki, M., Down-regulation of calcium/calmodulin-dependent protein kinase kinase 2 by androgen deprivation induces castration-resistant prostate cancer. 2012, 72, (16), 1789–1801.

81. Subbannayya, Y.; Syed, N.; Barbhuiya, M. A.; Raja, R.; Marimuthu, A.; Sahasrabuddhe, N.; Pinto, S. M.; Manda, S. S.; Renuse, S.; Manju, H. C.; Zameer, M. A. L.; Sharma, J.; Brait, M.; Srikumar, K.; Roa, J. C.; Vijaya Kumar, M.; Kumar, K. V. V.; Prasad, T. S. K.; Ramaswamy, G.; Kumar, R. V.; Pandey, A.; Gowda, H.; Chatterjee, A., Calcium calmodulin dependent kinase kinase 2 - a novel therapeutic target for gastric adenocarcinoma. Cancer Biology & Therapy 2015, 16, (2), 336–345.

82. Tan, M. H.; Li, J.; Xu, H. E.; Melcher, K.; Yong, E. L., Androgen receptor: structure, role in prostate cancer and drug discovery. Acta Pharmacol Sin 2015, 36, (1), 3-23.*

83. Lonergan, P. E.; Tindall, D. J., Androgen receptor signaling in prostate cancer development and progression. J Carcinog 2011, 10, (1), 20.

84. Frigo, D. E.; Howe, M. K.; Wittmann, B. M.; Brunner, A. M.; Cushman, I.; Wang, Q.; Brown, M.; Means, A. R.; McDonnell, D. P., CaM Kinase Kinase β-Mediated Activation of the Growth Regulatory Kinase AMPK Is Required for Androgen-Dependent Migration of Prostate Cancer Cells. 2011, 71, (2), 528–537.

85. Fu, H.; He, H.-c.; Han, Z.-d.; Wan, Y.-p.; Luo, H.-w.; Huang, Y.-q.; Cai, C.; Liang, Y.-x.; Dai, Q.-s.; Jiang, F.-n.; Zhong, W.-d. J. T. B., MicroRNA-224 and its target CAMKK2 synergistically influence tumor progression and patient prognosis in prostate cancer. 2015, 36, (3), 1983–1991.

86. Yasui, K.; Hashimoto, E.; Komorizono, Y.; Koike, K.; Arii, S.; Imai, Y.; Shima, T.; Kanbara, Y.; Saibara, T.; Mori, T.; Kawata, S.; Uto, H.; Takami, S.; Sumida, Y.; Takamura, T.; Kawanaka, M.; Okanoue, T.; Japan Nash Study Group, M. o. H. L.; Welfare of, J., Characteristics of patients with nonalcoholic steatohepatitis who develop hepatocellular carcinoma. Clin Gastroenterol Hepatol 2011, 9, (5), 428-33; quiz e50.

87. Takuma, Y.; Nouso, K., Nonalcoholic steatohepatitis-associated hepatocellular carcinoma: our case series and literature review. World J Gastroenterol 2010, 16, (12), 1436–41.

88. Anderson, K. A.; Ribar, T. J.; Lin, F.; Noeldner, P. K.; Green, M. F.; Muehlbauer, M. J.; Witters, L. A.; Kemp, B. E.; Means, A. R., Hypothalamic CaMKK2 contributes to the regulation of energy balance. Cell Metab 2008, 7, (5), 377–88.

89. Price, D. J.; Drewry, D. H.; Schaller, L. T.; Thompson, B. D.; Reid, P. R.; Maloney, P. R.; Liang, X.; Banker, P.; Buckholz, R. G.; Selley, P. K.; McDonald, O. B.; Smith, J. L.; Shearer, T. W.; Cox, R. F.; Williams, S. P.; Reid, R. A.; Tacconi, S.; Faggioni, F.; Piubelli, C.; Sartori, I.; Tessari, M.; Wang, T. Y., An orally available, brain-penetrant CAMKK2 inhibitor reduces food intake in rodent model. Bioorganic & Medicinal Chemistry Letters 2018, 28, (10), 1958–1963.

90. Anderson, K. A.; Ribar, T. J.; Lin, F.; Noeldner, P. K.; Green, M. F.; Muehlbauer, M. J.; Witters, L. A.; Kemp, B. E.; Means, A. R., Hypothalamic CaMKK2 Contributes to the Regulation of Energy Balance. Cell Metabolism 2008, 7, (5), 377–388.

91. York, B.; Li, F.; Lin, F.; Marcelo, K. L.; Mao, J.; Dean, A.; Gonzales, N.; Gooden, D.; Maity, S.; Coarfa, C.; Putluri, N.; Means, A. R., Pharmacological inhibition of CaMKK2 with the selective antagonist STO-609 regresses NAFLD. Sci Rep 2017, 7, (1), 11793.

92. Lin, F.; Marcelo, K. L.; Rajapakshe, K.; Coarfa, C.; Dean, A.; Wilganowski, N.; Robinson, H.; Sevick, E.; Bissig, K.-D.; Goldie, L. C.; Means, A. R.; York, B., The camKK2/camKIV relay is an essential regulator of hepatic cancer. Hepatology 2015, 62, (2), 505–520.

93. Ma, Z.; Wen, D.; Wang, X.; Yang, L.; Liu, T.; Liu, J.; Zhu, J.; Fang, X., Growth inhibition of human gastric adenocarcinoma cells in vitro by STO-609 is independent of calcium/calmodulin-dependent protein kinase kinase-beta and adenosine monophosphate-activated protein kinase. American journal of translational research 2016, 8, (2), 1164–1171.

94. Racioppi, L.; Nelson, E. R.; Huang, W.; Mukherjee, D.; Lawrence, S. A.; Lento, W.; Masci, A. M.; Jiao, Y.; Park, S.; York, B.; Liu, Y.; Baek, A. E.; Drewry, D. H.; Zuercher, W. J.; Bertani, F. R.; Businaro, L.; Geradts, J.; Hall, A.; Means, A. R.; Chao, N.; Chang, C. Y.; McDonnell, D. P., CaMKK2 in myeloid cells is a key regulator of the immune-suppressive microenvironment in breast cancer. Nat Commun 2019, 10, (1), 2450.

95. Racioppi, L.; Nelson, E. R.; Huang, W.; Mukherjee, D.; Lawrence, S. A.; Lento, W.; Masci, A. M.; Jiao, Y.; Park, S.; York, B.; Liu, Y.; Baek, A. E.; Drewry, D. H.; Zuercher, W. J.; Bertani, F. R.; Businaro, L.; Geradts, J.; Hall, A.; Means, A. R.; Chao, N.; Chang, C.-y.; McDonnell, D. P., CaMKK2 in myeloid cells is a key regulator of the immune-suppressive microenvironment in breast cancer. Nature Communications 2019, 10, (1), 2450.

96. Davare, M. A.; Saneyoshi, T.; Soderling, T. R. J. J. o. N.-O., Calmodulin-kinases regulate basal and estrogen stimulated medulloblastoma migration via Rac1. 2011, 104, (1), 65–82.

97. Cary, R. L.; Waddell, S.; Racioppi, L.; Long, F.; Novack, D. V.; Voor, M. J.; Sankar, U., Inhibition of Ca(2)(+)/calmodulin-dependent protein kinase kinase 2 stimulates osteoblast formation and inhibits osteoclast differentiation. J Bone Miner Res 2013, 28, (7), 1599–610.

98. Cary, R. L.; Waddell, S.; Racioppi, L.; Long, F.; Novack, D. V.; Voor, M. J.; Sankar, U., Inhibition of Ca2+/Calmodulin–Dependent Protein Kinase Kinase 2 Stimulates Osteoblast Formation and Inhibits Osteoclast Differentiation. Journal of Bone and Mineral Research 2013, 28, (7), 1599–1610.

99. Pritchard, Z. J.; Cary, R. L.; Yang, C.; Novack, D. V.; Voor, M. J.; Sankar, U., Inhibition of CaMKK2 reverses age-associated decline in bone mass. Bone 2015, 75, 120–7.

100. Lin, F.; Ribar, T. J.; Means, A. R., The Ca2+/calmodulin-dependent protein kinase kinase, CaMKK2, inhibits preadipocyte differentiation. Endocrinology 2011, 152, (10), 3668–79.

101. Kang, X.; Cui, C.; Wang, C.; Wu, G.; Chen, H.; Lu, Z.; Chen, X.; Wang, L.; Huang, J.; Geng, H.; Zhao, M.; Chen, Z.; Muschen, M.; Wang, H. Y.; Zhang, C. C., CAMKs support development of acute myeloid leukemia. J Hematol Oncol 2018, 11, (1), 30.

102. Pritchard, Z. J.; Cary, R. L.; Yang, C.; Novack, D. V.; Voor, M. J.; Sankar, U., Inhibition of CaMKK2 reverses age-associated decline in bone mass. Bone 2015, 75, 120–127.

103. Kukimoto-Niino, M.; Yoshikawa, S.; Takagi, T.; Ohsawa, N.; Tomabechi, Y.; Terada, T.; Shirouzu, M.; Suzuki, A.; Lee, S.; Yamauchi, T.; Okada-Iwabu, M.; Iwabu, M.; Kadowaki, T.; Minokoshi, Y.; Yokoyama, S., Crystal structure of the CA2+/calmodulin-dependent protein kinase kinase in complex with the inhibitor STO-609. Journal of Biological Chemistry 2011.

104. York, B.; Li, F.; Lin, F.; Marcelo, K. L.; Mao, J.; Dean, A.; Gonzales, N.; Gooden, D.; Maity, S.; Coarfa, C.; Putluri, N.; Means, A. R., Pharmacological inhibition of CaMKK2 with the selective antagonist STO-609 regresses NAFLD. Scientific Reports 2017, 7, (1), 11793.

105. Ritchie, T. J.; Macdonald, S. J., The impact of aromatic ring count on compound developability--are too many aromatic rings a liability in drug design? Drug Discov Today 2009, 14, (21-22), 1011–20.

106. Ritchie, T. J.; Macdonald, S. J.; Young, R. J.; Pickett, S. D., The impact of aromatic ring count on compound developability: further insights by examining carbo- and hetero-aromatic and -aliphatic ring types. Drug Discov Today 2011, 16, (3-4), 164-71.

107. Timm, M.; Saaby, L.; Moesby, L.; Hansen, E. W., Considerations regarding use of solvents in in vitro cell based assays. Cytotechnology 2013, 65, (5), 887–94.

108. Bain, J.; Plater, L.; Elliott, M.; Shpiro, N.; Hastie, C. J.; McLauchlan, H.; Klevernic, I.; Arthur, J. S.; Alessi, D. R.; Cohen, P., The selectivity of protein kinase inhibitors: a further update. Biochem J 2007, 408, (3), 297–315.

109. Morishita, D.; Katayama, R.; Sekimizu, K.; Tsuruo, T.; Fujita, N., Pim kinases promote cell cycle progression by phosphorylating and down-regulating p27Kip1 at the transcriptional and posttranscriptional levels. Cancer Res 2008, 68, (13), 5076–85.

110. Fujii, C.; Nakamoto, Y.; Lu, P.; Tsuneyama, K.; Popivanova, B. K.; Kaneko, S.; Mukaida, N., Aberrant expression of serine/threonine kinase Pim-3 in hepatocellular carcinoma development and its role in the proliferation of human hepatoma cell lines. Int J Cancer 2005, 114, (2), 209–18.

111. Li, Y. Y.; Popivanova, B. K.; Nagai, Y.; Ishikura, H.; Fujii, C.; Mukaida, N., Pim-3, a proto-oncogene with serine/threonine kinase activity, is aberrantly expressed in human pancreatic cancer and phosphorylates bad to block bad-mediated apoptosis in human pancreatic cancer cell lines. Cancer Res 2006, 66, (13), 6741–7.

112. Popivanova, B. K.; Li, Y. Y.; Zheng, H.; Omura, K.; Fujii, C.; Tsuneyama, K.; Mukaida, N., Proto-oncogene, Pim-3 with serine/threonine kinase activity, is aberrantly expressed in human colon cancer cells and can prevent Bad-mediated apoptosis. Cancer Sci 2007, 98, (3), 321–8.

113. Zheng, H. C.; Tsuneyama, K.; Takahashi, H.; Miwa, S.; Sugiyama, T.; Popivanova, B. K.; Fujii, C.; Nomoto, K.; Mukaida, N.; Takano, Y., Aberrant Pim-3 expression is involved in gastric adenoma-adenocarcinoma sequence and cancer progression. J Cancer Res Clin Oncol 2008, 134, (4), 481–8.

114. Monteiro, P.; Gilot, D.; Langouet, S.; Fardel, O., Activation of the aryl hydrocarbon receptor by the calcium/calmodulin-dependent protein kinase kinase inhibitor 7-oxo-7H-benzimidazo[2,1-a]benz[de]isoquinoline-3-carboxylic acid (STO-609). Drug Metab Dispos 2008, 36, (12), 2556–63.

115. Bain, J.; Plater, L.; Elliott, M.; Shpiro, N.; Hastie, C. J.; McLauchlan, H.; Klevernic, I.; Arthur, J. S. C.; Alessi, D. R.; Cohen, P., The selectivity of protein kinase inhibitors: a further update. Biochem J 2007, 408, (3), 297–315.

116. Landesman-Bollag, E.; Romieu-Mourez, R.; Song, D. H.; Sonenshein, G. E.; Cardiff, R. D.; Seldin, D. C., Protein kinase CK2 in mammary gland tumorigenesis. Oncogene 2001, 20, (25), 3247–57.

117. Daya-Makin, M.; Sanghera, J. S.; Mogentale, T. L.; Lipp, M.; Parchomchuk, J.; Hogg, J. C.; Pelech, S. L., Activation of a Tumor-associated Protein Kinase (p40<sup>TAK</sup>) and Casein Kinase 2 in Human Squamous Cell Carcinomas and Adenocarcinomas of the Lung. Cancer Research 1994, 54, (8), 2262–2268.

118. Yenice, S.; Davis, A. T.; Goueli, S. A.; Akdas, A.; Limas, C.; Ahmed, K., Nuclear casein kinase 2 (CK-2) activity in human normal, benign hyperplastic, and cancerous prostate. Prostate 1994, 24, (1), 11–6.

119. Stalter, G.; Siemer, S.; Becht, E.; Ziegler, M.; Remberger, K.; Issinger, O. G., Asymmetric expression of protein kinase CK2 subunits in human kidney tumors. Biochem Biophys Res Commun 1994, 202, (1), 141–7.

120. Duncan, J. S.; Litchfield, D. W., Too much of a good thing: the role of protein kinase CK2 in tumorigenesis and prospects for therapeutic inhibition of CK2. Biochim Biophys Acta 2008, 1784, (1), 33–47.

121. Li, L.; Liu, C.; Amato, R. J.; Chang, J. T.; Du, G.; Li, W., CDKL2 promotes epithelial-mesenchymal transition and breast cancer progression. Oncotarget 2014, 5, (21), 10840–53.

122. Billard, M. J.; Fitzhugh, D. J.; Parker, J. S.; Brozowski, J. M.; McGinnis, M. W.; Timoshchenko, R. G.; Serafin, D. S.; Lininger, R.; Klauber-Demore, N.; Sahagian, G.; Truong, Y. K.; Sassano, M. F.; Serody, J. S.; Tarrant, T. K., G Protein Coupled Receptor Kinase 3 Regulates Breast Cancer Migration, Invasion, and Metastasis. PLoS One 2016, 11, (4), e0152856.

123. Maloveryan, A.; Finta, C.; Osterlund, T.; Kogerman, P., A possible role of mouse Fused (STK36) in Hedgehog signaling and Gli transcription factor regulation. J Cell Commun Signal 2007, 1, (3-4), 165-73.

124. Arrowsmith, C. H.; Audia, J. E.; Austin, C.; Baell, J.; Bennett, J.; Blagg, J.; Bountra, C.; Brennan, P. E.; Brown, P. J.; Bunnage, M. E.; Buser-Doepner, C.; Campbell, R. M.; Carter, A. J.; Cohen, P.; Copeland, R. A.; Cravatt, B.; Dahlin, J. L.; Dhanak, D.; Edwards, A. M.; Frederiksen, M.; Frye, S. V.; Gray, N.; Grimshaw, C. E.; Hepworth, D.; Howe, T.; Huber, K. V.; Jin, J.; Knapp, S.; Kotz, J. D.; Kruger, R. G.; Lowe, D.; Mader, M. M.; Marsden, B.; Mueller-Fahrnow, A.; Muller, S.; O’Hagan, R. C.; Overington, J. P.; Owen, D. R.; Rosenberg, S. H.; Roth, B.; Ross, R.; Schapira, M.; Schreiber, S. L.; Shoichet, B.; Sundstrom, M.; Superti-Furga, G.; Taunton, J.; Toledo-Sherman, L.; Walpole, C.; Walters, M. A.; Willson, T. M.; Workman, P.; Young, R. N.; Zuercher, W. J., The promise and peril of chemical probes. Nat Chem Biol 2015, 11, (8), 536–41.

125. Blagg, J.; Workman, P., Choose and Use Your Chemical Probe Wisely to Explore Cancer Biology. Cancer Cell 2017, 32, (1), 9–25.

126. Asquith, R. C.; Godoi, H. P.; Couñago, M. R.; Laitinen, T.; Scott, W. J.; Langendorf, G. C.; Oakhill, S. J.; Drewry, H. D.; Zuercher, J. W.; Koutentis, A. P.; Willson, M. T.; Kalogirou, S. A., 1,2,6-Thiadiazinones as Novel Narrow Spectrum Calcium/Calmodulin-Dependent Protein Kinase Kinase 2 (CaMKK2) Inhibitors. Molecules 2018, 23, (5).

127. Ghose, A. K.; Herbertz, T.; Pippin, D. A.; Salvino, J. M.; Mallamo, J. P., Knowledge based prediction of ligand binding modes and rational inhibitor design for kinase drug discovery. J Med Chem 2008, 51, (17), 5149–71.

128. Krishna, S. N.; Luan, C. H.; Mishra, R. K.; Xu, L.; Scheidt, K. A.; Anderson, W. F.; Bergan, R. C., A fluorescence-based thermal shift assay identifies inhibitors of mitogen activated protein kinase kinase 4. PLoS One 2013, 8, (12), e81504.

129. Simeonov, A., Recent developments in the use of differential scanning fluorometry in protein and small molecule discovery and characterization. Expert Opin Drug Discov 2013, 8, (9), 1071–82.

130. Profeta, G. S.; Dos Reis, C. V.; Santiago, A. D. S.; Godoi, P. H. C.; Fala, A. M.; Wells, C. I.; Sartori, R.; Salmazo, A. P. T.; Ramos, P. Z.; Massirer, K. B.; Elkins, J. M.; Drewry, D. H.; Gileadi, O.; Counago, R. M., Binding and structural analyses of potent inhibitors of the human Ca(2+)/calmodulin dependent protein kinase kinase 2 (CAMKK2) identified from a collection of commercially-available kinase inhibitors. Sci Rep 2019, 9, (1), 16452.

131. Profeta, G. S.; dos Reis, C. V.; Santiago, A. d. S.; Godoi, P. H. C.; Fala, A. M.; Wells, C. I.; Sartori, R.; Salmazo, A. P. T.; Ramos, P. Z.; Massirer, K. B.; Elkins, J. M.; Drewry, D. H.; Gileadi, O.; Couñago, R. M., Binding and structural analyses of potent inhibitors of the human Ca2+/calmodulin dependent protein kinase kinase 2 (CAMKK2) identified from a collection of commercially-available kinase inhibitors. Scientific Reports 2019, 9, (1), 16452.

132. Zhang, S.; Anjum, R.; Squillace, R.; Nadworny, S.; Zhou, T.; Keats, J.; Ning, Y.; Wardwell, S. D.; Miller, D.; Song, Y.; Eichinger, L.; Moran, L.; Huang, W. S.; Liu, S.; Zou, D.; Wang, Y.; Mohemmad, Q.; Jang, H. G.; Ye, E.; Narasimhan, N.; Wang, F.; Miret, J.; Zhu, X.; Clackson, T.; Dalgarno, D.; Shakespeare, W. C.; Rivera, V. M., The Potent ALK Inhibitor Brigatinib (AP26113) Overcomes Mechanisms of Resistance to First- and Second-Generation ALK Inhibitors in Preclinical Models. Clin Cancer Res 2016, 22, (22), 5527–5538.

133. Davis, M. I.; Hunt, J. P.; Herrgard, S.; Ciceri, P.; Wodicka, L. M.; Pallares, G.; Hocker, M.; Treiber, D. K.; Zarrinkar, P. P., Comprehensive analysis of kinase inhibitor selectivity. Nat Biotechnol 2011, 29, (11), 1046–51.

134. Gohda, J.; Suzuki, K.; Liu, K.; Xie, X.; Takeuchi, H.; Inoue, J. I.; Kawaguchi, Y.; Ishida, T., BI-2536 and BI-6727, dual Polo-like kinase/bromodomain inhibitors, effectively reactivate latent HIV-1. Sci Rep 2018, 8, (1), 3521.

135. Zhang, S.; Anjum, R.; Squillace, R.; Nadworny, S.; Zhou, T.; Keats, J.; Ning, Y.; Wardwell, S. D.; Miller, D.; Song, Y.; Eichinger, L.; Moran, L.; Huang, W.-S.; Liu, S.; Zou, D.; Wang, Y.; Mohemmad, Q.; Jang, H. G.; Ye, E.; Narasimhan, N.; Wang, F.; Miret, J.; Zhu, X.; Clackson, T.; Dalgarno, D.; Shakespeare, W. C.; Rivera, V. M., The Potent ALK Inhibitor Brigatinib (AP26113) Overcomes Mechanisms of Resistance to First- and Second-Generation ALK Inhibitors in Preclinical Models. Clinical Cancer Research 2016, 22, (22), 5527.

136. Xing, L.; Klug-Mcleod, J.; Rai, B.; Lunney, E. A., Kinase hinge binding scaffolds and their hydrogen bond patterns. Bioorg Med Chem 2015, 23, (19), 6520–7.

137. Fedorov, O.; Niesen, F. H.; Knapp, S., Kinase Inhibitor Selectivity Profiling Using Differential Scanning Fluorimetry. In Kinase Inhibitors: Methods and Protocols, Kuster, B., Ed. Humana Press: Totowa, NJ, 2012; pp 109–118.

138. Niesen, F. H.; Berglund, H.; Vedadi, M., The use of differential scanning fluorimetry to detect ligand interactions that promote protein stability. Nat Protoc 2007, 2, (9), 2212–21.

