## Supplementary material for "In depth analysis of kinase cross screening data to identify CAMKK2 inhibitory scaffolds": SI

Supplementary Table S1: KINOME*scan* data for STO-609

Supplementary Table S2: Names and SMILES strings of compounds tested in this manuscript

Supplementary Table S3: Reaction Biology Corporation enzyme profiling data for GSK650394

Supplementary Methods:

Databases utilized

'Kinase Profiling Inhibitor Database' provided by the International Center for Kinase profiling within the MRC Protein Phosphorylation Unit at the University of Dundee.

http://www.kinase-screen.mrc.ac.uk/kinase-inhibitors

'Kinase Inhibitor Resource' provided by the Fox Chase Cancer Center and Reaction Biology,

http://reactionbiology.com/webapps/largedata/

'KInhibition' provided by Fred Hutch Cancer Research Center

https://kinhibition.fredhutch.org

LINCS KINOME*scan* database provided by Harvard Medical School.

http://lincs.hms.harvard.edu/kinomescan/
